# Causes and consequences of facultative sea crossing in a soaring migrant

**DOI:** 10.1101/758102

**Authors:** Paolo Becciu, Shay Rotics, Nir Horvitz, Michael Kaatz, Wolfgang Fiedler, Damaris Zurell, Andrea Flack, Florian Jeltsch, Martin Wikelski, Ran Nathan, Nir Sapir

## Abstract

1. Studying the causes and consequences of route selection in animal migration is important for understanding the evolution of migratory systems and how they may be affected by environmental factors at various spatial and temporal scales. One key decision during migration is whether to cross “high transport cost” areas, or to circumvent them. Soaring birds may face this choice when encountering waterbodies where convective updrafts are weak or scarce. Crossing these waterbodies requires flying using energetically costly flapping flight, while circumventing them over land permits energetically cheap soaring.
2. We tested how several atmospheric factors (e.g., wind, thermal uplift) and geographic, seasonal and state-related factors (sex and age) affected route selection in migrating white storks (*Ciconia ciconia*). We used 196 GPS tracks of 70 individuals either crossing or circumventing the north-easternmost section of the Mediterranean Sea, over Iskenderun Bay in southern Turkey.
3. We found that westward and southward winds promoted a cross-bay journey in spring and autumn, respectively, acting as tailwinds. Also, overall weaker winds promoted a sea crossing in spring. Sea crossing was associated with flapping flight and higher values of Overall Dynamic Body Acceleration (ODBA) and resulted in higher ground speed than travel over land.
4. The combined environmental conditions and the effects of route selection on movement-related energy costs and speed were likely responsible for an increase in the time spent flying and distance travelled of migrating storks that decided to cross the bay during spring. Notably, daily travel distances of spring migrants crossing the bay were 60 kilometres longer than those of land-detouring birds, allowing them to reach their destination faster but likely incurring a higher energetic flight cost. No such benefit was found during autumn.
5. Our findings confirm that atmospheric conditions can strongly affect bird route selection. Consequently, migration timing, speed and movement-related energy expenditure differed considerably between the two migratory seasons and the two route choices, highlighting a time-energy trade-off in the migration of white storks.

## 1. Introduction

Environmental conditions during long-distance bird migration are known to affect migration timing, flight performance and energy expenditure (Becciu et al., 2019; Shamoun-Baranes, Liechti, & Vansteelant, 2017). Still, how migration route is influenced by atmospheric and geographical factors is much less clear. Route selection over ecological barriers such as large waterbodies may depend on weather and geographical features (Alerstam, 2001; Becciu et al., 2019; Efrat, Hatzofe, & Nathan, 2019; Eisaguirre et al., 2018; Nourani, Yamaguchi, Manda, & Higuchi, 2016), affecting migration time and energy expenditures, with consequences for animal fitness (Shamoun-Baranes, Bouten, & Van Loon, 2010; Shamoun-Baranes et al., 2017). Large terrestrial soaring birds depend on local atmospheric conditions during their flight, since they utilize thermal uplifts to gain height and later glide towards their destination (Norberg, 1990). During soaring flight, the birds stretch and do not flap their wings, allowing them to save energy while covering large distances (Sapir, Wikelski, Mccue, Pinshow, & Nathan, 2010). Usually, soaring birds avoid flying over waterbodies where thermals are typically weak and rare (but see Duriez, Peron, Gremillet, Sforzi, & Monti, 2018; Nourani, Vansteelant, Byholm, & Safi, 2019). Yet, in some cases soaring birds are forced to switch to the metabolically demanding flapping flight (Hedenström, 1993; Norberg, 1990; Sapir et al., 2011), such as when flying over areas with low availability of thermals. These areas can be regarded as “high transport cost” areas (Alerstam, 2001). We note that differences in transport cost may be caused by additional factors, such as variable wind conditions experienced by the birds when travelling over different areas (Alerstam, 2001; Efrat et al., 2019). Besides increasing the transport cost, barrier crossing versus barrier circumvention (i.e., facultative barrier crossing) may shorten migration distance, with possible consequences for migration time saving (Alerstam, 2001; Efrat et al., 2019).

Weather conditions may affect the timing and the location of the crossing in obligatory sea crossing during migration (Agostini, Panuccio, & Pasquaretta, 2015; Bildstein, 2006; Bildstein, Bechard, Farmer, & Newcomb, 2009; Meyer, Spaar, & Bruderer, 2000; Nourani et al., 2016). For example, Oriental honey-buzzards (*Pernis ptilorhynchus*) that crossed the sea between the mainland and Japan were affected by wind conditions and the geography of the study area (Nourani et al., 2016; Yamaguchi, Arisawa, Shimada, & Higuchi, 2012). Wind conditions also affected the propensity of several species of soaring migrants to cross the area of the strait of Gibraltar in locations where the cross-sea travel was not the shortest possible (Meyer et al., 2000). Compared to the latter situations of obligatory sea crossing, causes and consequences of a facultative sea-crossing decision in soaring migrants were rarely studied to date (Kerlinger, 1984).

We investigated the flight behaviour of the white stork, *Ciconia ciconia*, a large soaring bird and a long-distant migrant, as it passed through the Iskenderun Bay (“the bay” hereafter) in the north-eastern corner of the Mediterranean Sea. We found that about half of the birds crossed the bay over water while the other storks circumvented it over land. We examined how meteorological conditions affected migration route selection (bay crossing vs. overland detour) and furthermore explored the consequences of route selection for migration travel distance and movement-related energetics due to changes in the prevalence of the two flight modes (soaring vs. flapping) used by the birds. Large differences in flight energetic costs between the two flight modes (Sapir et al., 2010) imply a possible trade-off between different benefits and costs of facultative sea-crossing behaviour. For example, over-sea shortcutting may involve high prevalence of energetically expensive flapping flight whereas the longer overland detour might be associated with low-cost soaring flight. We consequently hypothesize that cross-sea flight is selected only when specific meteorological conditions prevail, such as tailwinds, which increase the benefit-(shorter travel time)-to-cost (high energetic costs due to flapping) ratio of crossing the bay. We therefore tested how wind speed and direction, temperature and thermal availability affected the decision of the storks to cross the bay. We furthermore considered the sex and the age of the individuals because intrinsic individual attributes may play an important role in determining movement decisions in general (Nathan et al., 2008), and specifically in migrating white storks (Rotics et al., 2016; Rotics et al., 2018). Also, we considered the timing of bird passage through the study area within the season. We additionally explored time and energy consequences of route selection. We hypothesize that sea-crossing behaviour is not random and depends on both extrinsic and intrinsic factors that could affect individual fitness. We specifically predict that: (a) tailwinds will facilitate sea-crossing flight, and increase the speed of migration (Becciu, Panuccio, Catoni, Dell’Omo, & Sapir, 2018; Nourani et al., 2016). (b) Early-spring migrants will show higher sea-crossing propensity due high motivation to arrive earlier at their breeding grounds, and more so in males (Rotics et al., 2018). Further, we expect juveniles which are less prone to risk to travel through a safer land detour (Harel et al., 2016; Rotics et al., 2016). Early-spring migrants may further show higher sea-crossing propensity due to poor thermal conditions over land in early spring (Rotics et al., 2018; Shamoun-Baranes et al., 2003). (c) Sea-crossing flight will be beneficial to the migrants, shortening their route distance and time compared with land detour, consequently allowing them to allocate the saved time to cover more distance at the end of the migration day. (d) Sea-crossing flight will require flapping as opposed to soaring during land detour and consequently will be metabolically more costly (Sapir et al., 2010; Wilson et al., 2019). (e) Sea crossing will not be the outcome of individual consistency in route choice over the years, which is a strategy that might have developed with experience or with individual preference (Vardanis, Klaassen, Strandberg, & Alerstam, 2011; Vardanis, Nilsson, Klaassen, Strandberg, & Alerstam, 2016), but rather mainly depend on local meteorological conditions before deciding whether to cross the sea. Therefore, we suggest that facultative sea-crossing behaviour could be the outcome of a time-energy trade-off during white stork migration, in which the birds may trade off energy expenditure for migration speed.

## 2. Materials and Methods

### 2.1 Bird tagging and study area

The white stork is a large long-distance migrant that breeds mainly in Europe and Western Asia, and the majority of its population over-winters in sub-Saharan Africa. The study took place at the area of Iskenderun Bay, Turkey (36.6330°N, 35.8786°E). White storks that migrate along the eastern Mediterranean flyway pass regularly over the study area twice a year. When encountering the bay, storks may choose to cross the bay, which is 30-45 km wide, or to circumvent it over land (Figure 1). From 2011 to 2014, we fitted solar-charged GPS transmitters with tri-axial acceleration (ACC) sensors (e-obs GmbH, Munich, Germany) to 62 adult and 84 immature white storks in the state of Saxony-Anhalt, Germany (see Rotics et al., 2016, 2017 for detailed methods regarding tagging and trapping protocols). Eight immature storks (birds in their first, second and third year of life) survived to the following years, allowing us to assess whether their behaviour changed with age. We found that the behaviour of 1^st^-year birds was similar to that of 2^nd^-and 3^rd^-year birds, in terms of sea crossing choice and day of passage over the study area (Figure S2), and consequently considered them in the same age class (juvenile) in the statistical analysis. Bird sex was determined by molecular methods (Rotics et al., 2018). The transmitters recorded GPS fixes every 5 minutes when solar conditions were good (95% of the time) or otherwise every 20 minutes. Every five minutes an ACC burst of 3.8 seconds was recorded at 10.54 Hz. Data were stored on-board and were downloaded via a VHF radio link upon locating the stork (Rotics et al., 2016). We excluded from the analysis tracks that did not present a clear route choice (storks that mostly followed the cost and cross less than 20 km over the bay), birds wintering at higher latitudes in the northern hemisphere (Rotics et al., 2017) and storks that did not cross the bay in one day (e.g. stopping over at the area of the bay). Consequently, we used data from 70 storks (39 adults and 31 immatures) that provided a total of 196 tracks (153 from adult and 43 from immature storks, 83 for spring and 113 for autumn migration). The maximum range of the storks’ tracks that travelled through the Iskenderun Bay area during a single day defined the geographic boundaries of the study, which were approximately 33.004° (westernmost longitude), 37.722° (easternmost longitude), 37.998° (northernmost latitude), and 34.963° (southernmost latitude).

**Figure 1.**
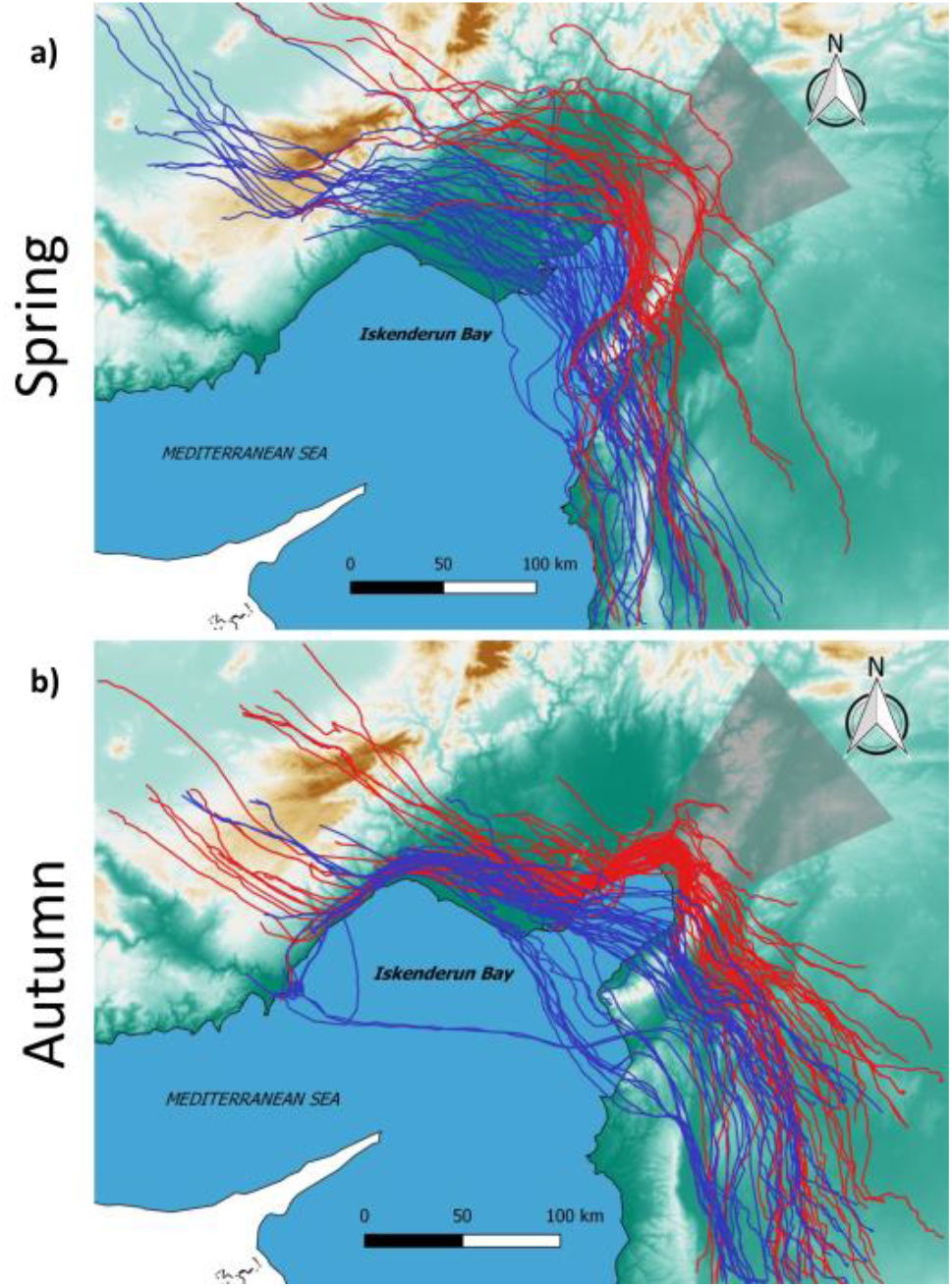
Map of the study area showing the white storks’ routes in (a) spring and (b) autumn. Blue routes depict sea crossing and red ones represent land detour. The shaded area is the bay-crossing stage named “ACROSS”, which is considered in the analysis of land-detouring birds (see Methods for details). The topography is depicted by a colour gradient from sea level (dark green) to mountains of about 3000 m a.s.l. (dark brown).

### 2.2 Movement parameters

Information regarding environmental data annotation of the birds’ tracks is provided in the supporting online material (S1). We calculated ground speed (*v_g_*) based on the time interval between two consecutive locations and additionally calculated the angle (*σ_i_*) of each such segment relative to the previous segment. These parameters were calculated using the package “move” in R (Kranstauber, Smolla, & Scharf, 2018). Ground speed was subsequently averaged for the entire day during which the bay crossing took place. For every ACC burst we calculated the birds’ Overall Dynamic Body Acceleration (ODBA), a valid proxy for activity-related energy expenditure (Wilson et al., 2019), and their flight mode (either flapping or soaring-gliding flight; see (Rotics et al., 2018) for details). Flight mode was annotated to each location and the proportion of flapping flight out of the total was calculated (proportion of gliding was one minus proportion of flapping) for a pre-defined area or for the birds’ daily travel over the area (see below the division of subsets). ODBA and the proportion of flapping flight are highly correlated (Spearman-ρ = 0.92, p < 0.001). Flight height above ground was calculated by subtracting ground elevation (obtained from ASTER ASTGTM2 Global 30-m DEM data set) (Dodge et al., 2013) and geoid height (the elevation difference between ellipsoid and geoid earth models) from the ellipsoid height recorded by the GPS transmitter. Air speed (*v*_*a*_) was calculated for each segment of the individual tracks following Safi et al. (2013): 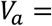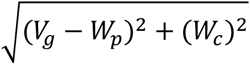.A single trip was defined from a starting point established where the ground speed exceeded 5 m/s after a nocturnal staging to an ending point where ground speed was below 2 m/s after a day of flight. We calculated time spent flying and distance travelled as cumulative sum of time and distance intervals at the day of the bay area crossing and at pre-defined sections of the daily trip (see below).

### 2.3 Data analysis

Our analyses were done considering tracks within a single day, during the time window when the storks were migrating (03:00 - 17:00 UTC). We divided our dataset into different subsets depending on the position of the birds with respect to the bay on the day of crossing the study area within the following three sections: 1) BEFORE (from take-off to the “bay area” – see below), 2) ACROSS (over the “bay area” or its projection over land), 3) AFTER (from the “bay area” until landing). A minimum of three consecutive locations per section was required for including data from a given section. The “bay area” is considered as the water body itself plus its projection over land in a direction perpendicular to the GPS tracks (shaded area in Figure 1, see also Figure S1). We used averaged movement and environmental data per day and per each bay-crossing section (depending on the analysis) to avoid spatial and temporal correlation on the day when the storks passed over the study area. We assigned bird tracks to two categories, namely LAND and SEA, for land-detour and sea-crossing routes, respectively.

To test the first part of prediction (a), as well as prediction (b), we tested bird route choice before arriving at the bay using Generalized Linear Mixed Models (hereafter GLMMs) with a binomial response variable (route choice: 0=LAND; 1=SEA), separately for autumn and spring migration, in relation to environmental factors, ordinal date and individual factors (i.e. sex, age) as well as two random factors (calendar year and bird ID). To avoid multicollinearity issues, we chose the most biologically meaningful variable from pairs of variables with a Spearman rank correlation | ρ | > 0.6. This ensured that all the predictors in the GLMMs had a Variance Inflation Factor (VIF) < 3 (Zuur, Ieno, & Elphick, 2010). We then tested all combinations of remaining variables in the global model and ranked the selected models according to the Akaike information criterion (Burnham & Anderson, 2002) using an automated stepwise model selection procedure in which models are fitted through repeated evaluation of modified calls extracted from the model containing all the meaningful variables, corrected for small sample sizes (AIC_c_) (Sugiura, 1978). Furthermore, we averaged all models with ∆AIC_c_ < 7 (Burnham, Anderson, & Huyvaert, 2011) and used the Akaike weights (*w*_*i*_) (Anderson, Burnham, & Thompson, 2000; Anderson, Link, Johnson, & Burnham, 2001) to assess the relative importance of the different variables. We used two global models, the first including E-W and N-S winds (but not *W*_*p*_ and *W*_*c*_), and the second with *W*_*p*_ and *W*_*c*_ (without E-W and N-S winds), and then used the one with the lowest AIC_c_ among them. We used 10-fold cross-validation with 10 repetitions, where the best model was trained on 70% of the data and then applied to the remaining 30% of the data. These data subsets were chosen randomly for each repetition (Hastie, Tibshirani, & Friedman, 2009; Meijer & Goeman, 2013). From the repeated cross-validation we reported the ability of the best model to distinguish between land/sea-crossing decisions using the area under the curve (AUC) of the receiver-operating characteristic curve (with standard deviation), the logistic regression accuracy (defined as the ratio between the sum of correct predicted cases of sea crossing and land detour and the sum of correct and non-correct predicted cases), sensitivity (proportion of land-detour choices correctly classified) and specificity (proportion of sea-crossing choices correctly classified) (Fawcett, 2006). To test prediction (e), individual consistency in route choice was examined by calculating repeatability across years (Intraclass correlation) using the rptR package (Stoffel, Nakagawa, & Schielzeth, 2017).

To test the second part of prediction (a) and prediction (c), we used linear mixed models (LMMs) to examine the effects of route choice and environmental factors on the daily ground and air speeds. We found the optimal structure of the fixed component as described above for GLMMs, using AIC_c_ in a multi-model selection framework. Also, we inspected GLMMs and LMMs residuals and considered the dispersion of the data (Zuur, 2009) using a simulation-based approach to create readily interpretable scaled (quantile) residuals for fitted (generalized) LMMs with the package DHARMa (Hartig, 2019). To test prediction (d) we additionally used LMMs to compare the two route choices in terms of time spent flying, distance covered, ground and air speeds, ODBA and proportion of flapping flight in of the daily travel and among the subsets (BEFORE, ACROSS and AFTER the bay). We report differences between route choices among the path segments using the *lsmeans()* R function of the package lsmeans (Lenth, 2016). Model fitting and multi-model inference were carried out in the statistical environment R 3.5.1 (R Core Team, 2018) by the packages lme4 (Bates, Mächler, Bolker, & Walker, 2015) and MuMIn (Barton, 2019), while the cross-validation was done using the package caret (Kuhn, 2019).

## 3. Results

### 3.1 Route selection

Migrating white storks crossed the Iskenderun Bay more often in spring (61.5%) than in autumn (39.8%). During the spring seasons of 2011 to 2015 storks crossed the Iskenderun Bay 51 times and detoured it 32 times. Adults preferred crossing the bay rather than detouring it (N_LAND_ = 24, N_SEA_ = 50; *χ*^2^ = 9.13, *p* < 0.01), while an opposite trend was found in juveniles (N_LAND_ = 8, N_SEA_ = 1). Juveniles travelled mostly over land also in autumn, (N_LAND_ = 25, N_SEA_ = 9; *χ*^2^ = 7.53, *p* < 0.01), whereas adults did not show any route selection preference in this season (N_LAND_ = 43, N_SEA_ = 36; *χ*^2^ = 0.62, *p* = 0.430).

In spring, wind speed, E-W wind speed and the ordinal date were ranked as the most important variables influencing route selection (Figure 2a) such that sea crossing was facilitated by decreasing wind speed, increasing westward wind speed and earlier passage date (Figure S12-14; Table S3-5). The average (± SD) logistic regression accuracy of the best-ranked model following the testing of the data subsets was 0.86 (± 0.13), with sensitivity_LAND_ = 0.76 (± 0.30) and specificity_SEA_ = 0.92 (± 0.14). The average (± SD) AUC was 0.96 (± 0.09). Route choice of individual birds was not consistent among years (n = 44; repeatability: *r* = 0.05 ± 0.09, *p* = 0.340).

**Figure 2.**
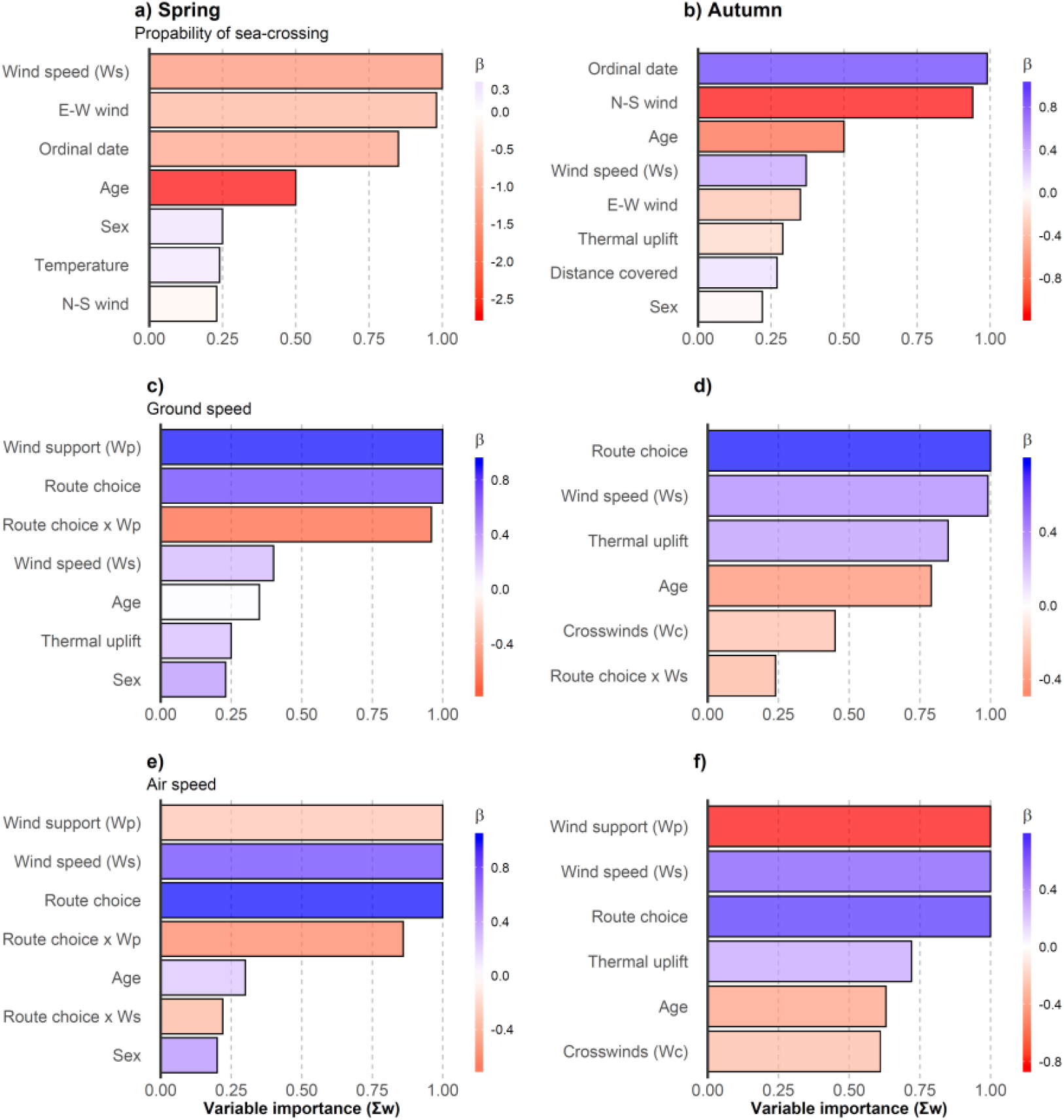
Summary of predictor averaged coefficients (*β*) ranked according to their predictive importance (Σ*w*) in models with ∆AIC_c_ < 7. Only results with a minimal Σ*w* = 0.2 are presented. Dependent variables are: probability of sea crossing (a, b), ground speed (c, d) and air speed (e, f). The baseline levels of the binomial variables “Route choice”, “Age” and “Sex” are land-detour (*LAND*), *adult* and *female*, respectively. See Tables S2-26 for a complete overview of the models’ procedure and results.

In autumn, N-S wind and ordinal date were the most influential factors affecting route selection (Figure 2b). The probability of sea crossing increased with southward wind speed and when passing over the area relatively late in the season (Figure S15-17; Table S8-9). The average (± SD) logistic regression accuracy of the best-ranked model was 0.74 (± 0.14), with sensitivity_LAND_ = 0.84 (± 0.17) and specificity_SEA_ = 0.60 (± 0.27). The average AUC was 0.85 (± 0.15). Also in autumn, route choice of individual birds was not consistent among years (n = 67; repeatability: *r* = 0.0001 ± 0.07, *p* = 0.626). Tables with model and variable rankings as well as the selected models are reported in the electronic supplementary material. In both seasons the best models with the lowest AIC_c_ values were those that included E-W and N-S winds and not *W_p_* and *W_c_* (ΔAIC_c_ = 3.41 in spring and 5.98 in autumn).

Figure 3 shows an overview of the winds available during the migration periods and the wind conditions (direction and speed) that the storks experienced before crossing or detouring the bay (BEFORE section; see also Figures S4-9).

**Figure 3.**
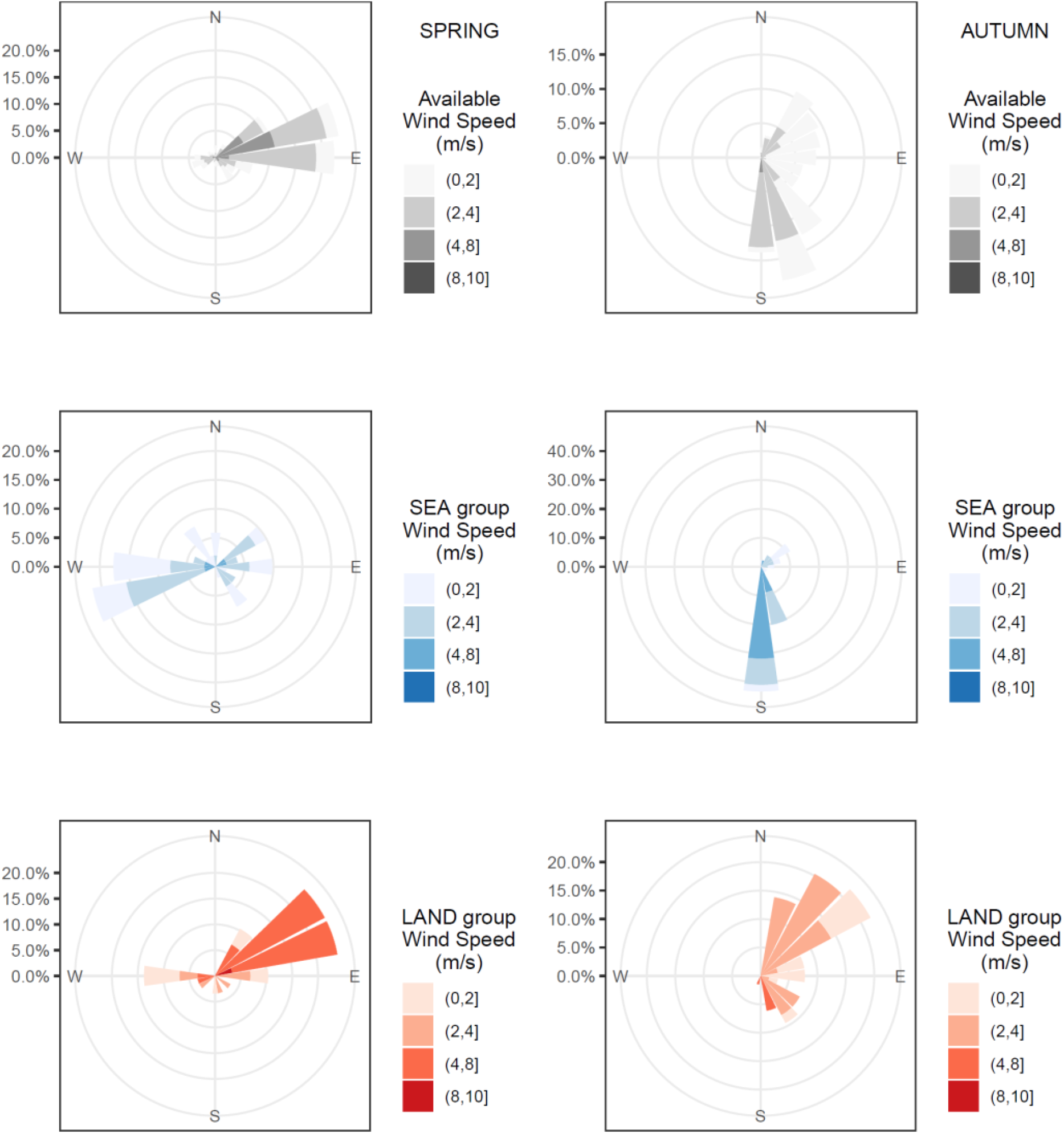
Wind roses plots of available winds and those used by white storks over the area BEFORE the bay. The available winds are depicted in grey tones, representing the daily averages of the entire period of migration window (± 2 days) for all the years of the study. The winds encountered by the storks before crossing the bay are depicted in blue tones, and those encountered by the birds that detoured the bay are illustrated in red tones. Plots on the left are from the spring, and those on the right are from the autumn. See also Figures S4-9.

### 3.2 Flight speed

#### 3.2.1 Ground speed

The storks’ ground speed was 7% higher on average in autumn than in spring (LMM: *β* = −0.77 ± 0.18, *t*_192_ = −4.3, *p* < 0.001). Considering their daily track and regardless of the season, they were 8% faster on average when crossing the sea than when flying over land (LMM: *β* = 0.7 ± 0.17, *t*_192_ = 3.99, *p* < 0.001). No difference in ground speed was found between adults and juveniles when data from the two migration seasons were pooled. In spring, white storks flew faster in tailwinds and slower under headwinds in general, but their route choice modulated their response (Figure 2c; Table S12-13). Over land, they increased their ground speed in tailwind (and decreased it under headwinds), but during sea crossing they maintained a rather steady ground speed regardless of wind support (Figure S3). In autumn, storks flew faster under stronger winds, thermal uplifts, crosswinds and when crossing the bay, compared to over-land flight. Also, adults flew faster than juveniles in this season (Figure 2d; Table S26). In spring, the best model with the lowest AIC_c_ value included *W_p_* and not E-W winds (ΔAIC_c_ = 11.45). In autumn, the two selected models (ΔAIC_c_ = 0) included either *W_p_* and *W_c_* or E-W and N-S winds.

#### 3.2.2 Airspeed

Overall, the storks’ daily airspeed was 7% higher on average in spring than in autumn (LMM: *β* = 0.55 ± 0.19, *t*_192_ = 2.89, *p* < 0.01), and adults were 9% faster on average than juveniles (LMM: *β* = −0.5 ± 0.22, *t*_192_ = −2.25, *p* = 0.025). Notably, considering data from both seasons, no significant difference in bird airspeed was found between detouring and bay-crossing storks. In spring, bay-crossing storks adjusted their airspeed to wind support (Figure 2e), decreasing it with tailwinds and increasing it with headwinds, while land detouring storks did not adjust their airspeed to wind conditions (Figure S3). Also, storks generally increased their airspeed with increasing wind speed (Figure 2e). In autumn, stork airspeed was higher under stronger headwinds, crosswinds and thermal uplifts and when crossing the bay (Figure 2f). In both seasons the best models with the lowest AIC_c_ values were those that included *W_p_* and *W_c_*, and not E-W and N-S winds (ΔAIC_c_ = 11.89 in spring and 40.81 in autumn).

### 3.3 Route choice and flight time, distance, energy and speed

We tested for differences in several flight parameters – namely time spent flying, distance covered, ODBA, proportion of flapping flight – between the two route choices (LAND or SEA) BEFORE, ACROSS and AFTER crossing the bay area, as well as over the entire daily path of the birds (Figure 3, Table S1). We found that the distance covered and the time spent flying depended on route choice. In spring, white storks that crossed the bay spent on average two more hours flying (see also Figures S20-23) and covered 60 km more distance, with the main difference found after crossing the bay, while in autumn the distance covered and the time spent flying were similar between the two route choices (Figure 3a,b). The average distance covered over the bay was 55.47 km (range: 28.12 – 144.33) in spring, and 70.50 km (range: 25.36 – 182.81) in autumn (Figure S10). ODBA and proportion of flapping flight were about 40% higher in storks that crossed the bay in both seasons (Figure 3c,d and Figure 4) in the day that included the cross-bay journey.

**Figure 4.**
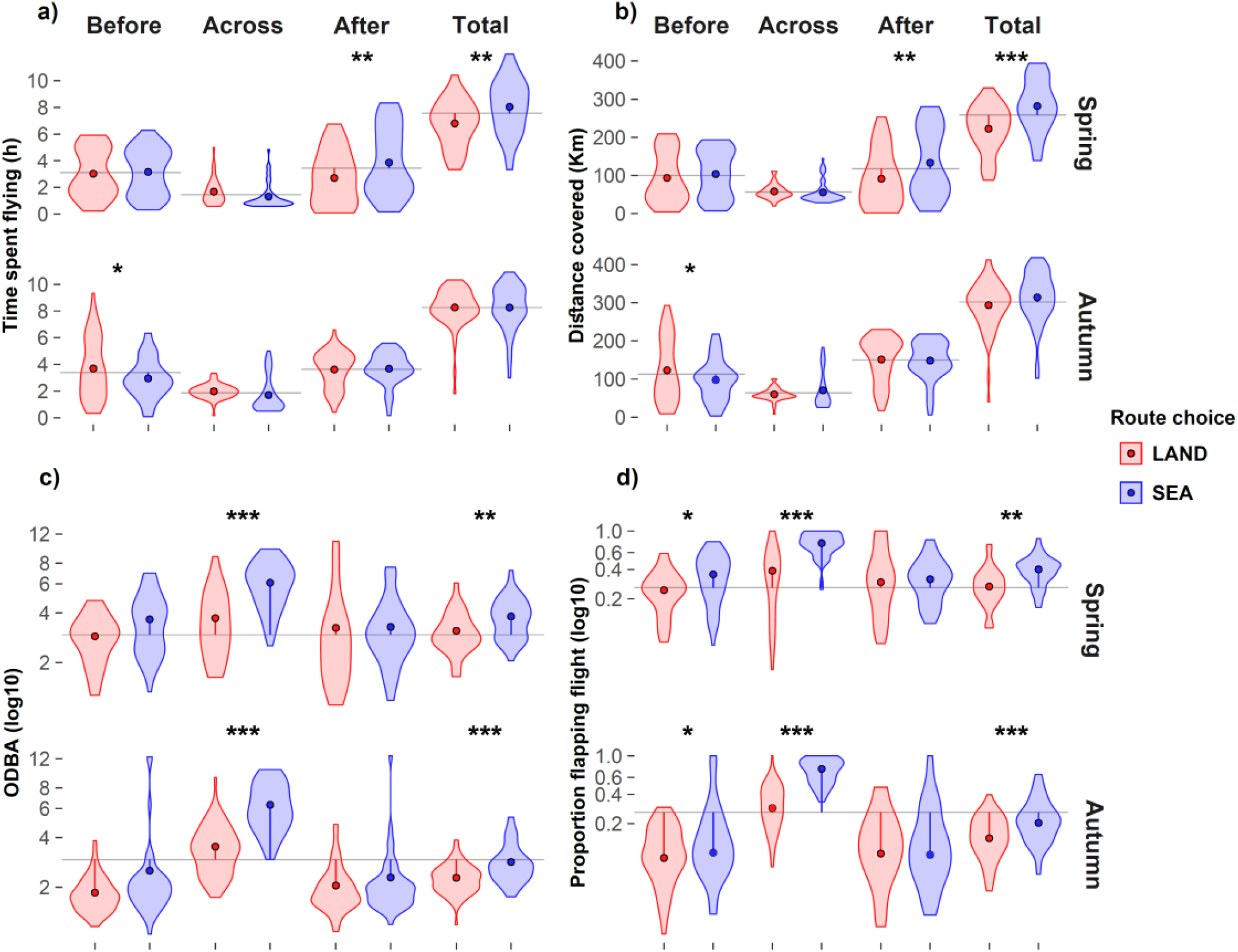
Summary statistics of (a) time spent flying, (b) distance covered, (c) Overall Dynamic Body Acceleration (ODBA), and (d) proportion of flapping flight of migrating storks flying over Iskenderun Bay area, according to the section of the flight path with respect to the bay (before, across or after) and the entire daily path, and by season. Colours represent the two route choices: land-detour (red) and sea-crossing (blue). Horizontal grey lines are averages per section and season (a, b) and overall average regardless of season and section (c, d). Dots are mean values and the shapes represent the distributions of the data. Asterisks indicate the *p*-value ranges: *p* < 0.001 (***), *p* < 0.01 (**), *p* < 0.05 (*). See also Figure S6 and S7 for explanations regarding differences in time and distance between the two route choices over the bay area.

**Figure 5.**
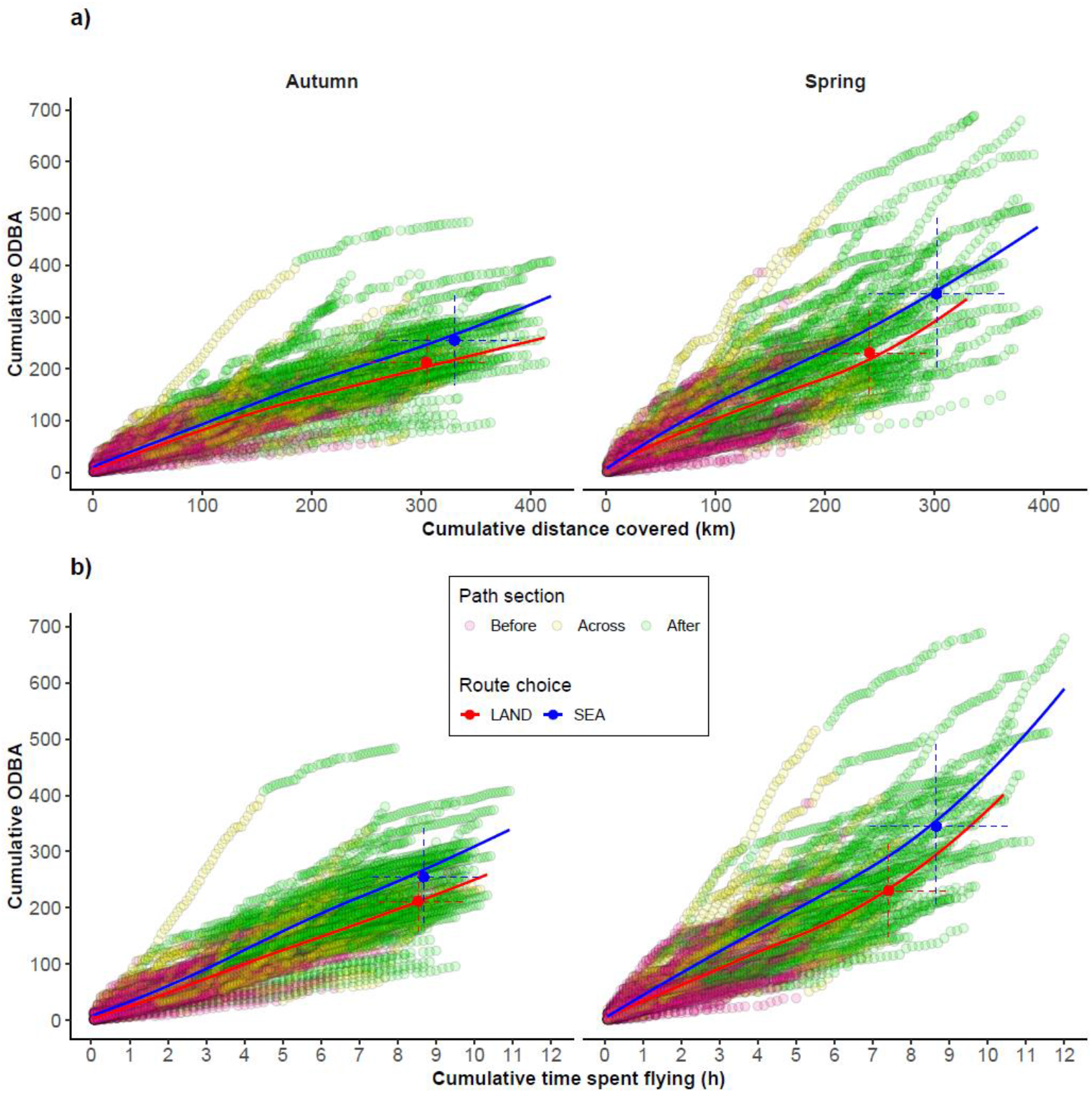
Visualization of cumulative sum of Overall Dynamic Body Acceleration (ODBA) in relation to cumulative distance covered (a) and cumulative time spent flying (b). The curves show the average relationships recorded from white storks that crossed the bay (blue) and those that circumvented it (red). Closed circles of these colours represent the mean (± SD) of each selected route choice (sea-crossing and over-land). Sequences of open coloured (see details below) circles depict data from white storks such that each sequence represents data from a single track across the bay area. The circles’ colour depicts the section over which they were recorded, with respect to the Iskenderun Bay: BEFORE (violet), ACROSS (yellow) and AFTER (green) the bay (see also electronic supplementary material, Figure S1).

## 4. Discussion

We highlight how important and consequential the choice of migration route is for soaring birds that either crossed a sea barrier or flew around it. Our findings uncover how migration route selection over a shorter path that is nonetheless characterized by a “high transport cost” is undertaken. In our case, the birds must flap over the sea, while flying a longer over-land detour route is associated with a lower transport cost because the birds are able to soar over it. We explored the factors that modulate route selection in a large soaring migrant, the white stork and inspected route selection consequences for flight behaviour, migration speed and flight energetics. Specifically, wind influenced route selection (Figure 3) which in turn affected bird ground and air speed, as well as the birds’ flight mode (soaring-gliding flight over land and flapping flight over the sea). Consequently, the combined effects of environmental conditions and route selection on energy costs and speed are likely responsible for the increase in the time spent flying and distance travelled of migrating storks that decided to cross the bay during spring. However, this longer daily migration distance came with a higher energetic flight cost, highlighting a likely time-energy trade-off in the migration of white storks. Yet, this benefit of sea crossing was found only in spring, allowing the birds to arrive earlier to their breeding grounds. The higher migratory motivation of those individuals that crossed the bay might have additionally played a major role in determining several aspects of their journey, including their daily travel duration. It is possible that the lower propensity of over-sea flights in autumn was based more on minimizing the risks during migration to reach the wintering grounds. Because route selection was strongly related to the local wind conditions at the day of passage, and was characterized by low repeatability, we hypothesize that route choice is not based on a fixed strategy of each individual but rather on a flexible selection with respect to local atmospheric conditions when arriving to the bay area. It is also noteworthy that the storks migrate in flocks, and thus route selection might not be an individual decision but rather a decision taken by the flock leaders (Flack, Nagy, Fiedler, Couzin, & Wikelski, 2018), possibly masking individual-related attributes. The lower rates of sea crossing in juveniles compared with adults could be related to their lower migratory experience (Rotics et al., 2016) and to lower migratory motivation since they do not breed. Possibly, juvenile birds trade off time and energy in a different manner than adults by responding more strongly to the negative aspects of the cross-bay flight.

Overall, we tested five predictions, how (a) tailwinds and (b) time pressure, sex and age could affect route choice, and how (c) sea-crossing could save time, possibly allowing extending the daily migration distance. We further tested whether storks have (d) higher energetic costs due to flapping flight while passing over the bay and whether (e) individual consistency played a role in bird route selection. Our first prediction (a) that tailwinds facilitate sea-crossing decision was confirmed. Decreasing wind speed, increasing westward winds in spring and increasing southward winds in autumn promoted sea crossing. The N-S and the E-W winds have a likely supporting effect in each season, since the crossing of the bay took place mostly from north to south in autumn and from east to west in spring. Similar results were reported by Meyer et al. (2000) for fall migration of soaring migrants crossing the Strait of Gibraltar with favourable southward and westward winds. In the same area Griffon vultures (*Gyps fulvus*) were also observed to cross the Strait of Gibraltar under weak winds or similarly with tailwinds (Bildstein et al., 2009).

Notably, we found contrasting responses to tailwinds between birds that selected the two routes in spring (see Figure S3). Specifically, birds that travelled over land increased their ground speed under tailwinds and decreased it under headwinds (see also Shamoun-Baranes et al., 2003), but kept a steady airspeed in both tailwind and headwind conditions. On the contrary, over the sea, when the birds employed flapping flight (see below), they adjusted their airspeed and maintained a quasi-steady ground speed, as observed in several studies of flapping birds and bats (Liechti, 1995; Sapir, Horvitz, Dechmann, Fahr, & Wikelski, 2014). No such differences in the birds’ response to the wind were found in autumn. We found a general increase in ground speed and decrease in airspeed with increasing tailwinds, suggesting that storks probably partially drifted with the wind in their preferred direction (over sea or over land). This is supported by the fact that the tracks were experiencing mostly tailwinds and almost no headwinds (Figure S3), meaning that they probably adjusted their movement to exploit those winds along the daily route in order to undertake a sea crossing or a land detour (Figure 3).

Our second prediction (b) was supported by our results since early-spring migrants commonly crossed the bay while relatively late migrants mostly detoured over land (Figure 2a and S14). We note that the higher tendency to cross the sea in spring and mostly with westward winds may be related to less suitable thermal conditions over land in spring that hindered soaring flight, compared to autumn (Figure S11). Furthermore, soaring conditions likely improved with ordinal date in spring, possibly explaining the increasing tendency for a land detour with the progression of the spring (Figure S11). Notably, early-spring migrants are typically considered as ‘higher-quality’ individuals, with better body condition (Dittmann & Becker, 2003), breeding success (Smith & Moore, 2005), and flight performance (Matyjasiak, 2013), which might explain their higher rates of selecting the shorter but energy demanding sea-crossing route.

Our findings partially support our third prediction (c) that sea-crossing flight is beneficial as it saves travelling time (see also Figures S19-23), and extends the daily distance travelled. Our data suggest that this was the case only for spring but not for autumn. The results support the prediction (d) that sea-crossing is associated with higher movement-related metabolic costs, since sea-crossing birds mostly used flapping flight and had higher ODBA (and thus likely higher flight energetic costs) compared with overland detouring birds. As predicted (e), no individual consistency was found in bird route selection.

In autumn, choosing one route or the other had no benefits in terms of more distance covered after the bay, but due to the use of flapping flight when crossing the bay, the birds that flew over the sea likely had higher flight energetic costs compared with land detouring birds that mostly flew using soaring flight. However, one has to bear in mind that since we found an effect of the prevailing meteorological conditions on route choice, we could not compare storks that used the two alternative routes under similar weather conditions. Hence, our comparison of migration performance between overland versus sea-crossing tracks are limited by the specific weather conditions that prevailed in the area in which the storks selected their route.

Understanding how atmospheric processes impact migration movements is of fundamental importance in a time of climate change (La Sorte, Horton, Nilsson, & Dokter, 2018; Nourani, Yamaguchi, & Higuchi, 2017; Winkler et al., 2014). In Turkey, including in the area of Iskenderun Bay, wind speed and specifically the E-W component of wind speed decreased over the last decades (Dadaser-Celik & Cengiz, 2014), partially following changes in global circulation patterns and increasing surface roughness (Vautard, Cattiaux, Yiou, Thépaut, & Ciais, 2010). Our results indicate that migrants are sensitive to the dynamics of their aerial environment and their behaviour and movement properties are strongly affected by local meteorological conditions. Changing atmospheric patterns due to climate change may thus result in changes in migration route selection of migrating white stroks, with possible implications for population dynamics (La Sorte, Fink, & Johnston, 2019) and conservation (Wilcove & Wikelski, 2008).

## Acknowledgements

We thank T. Schaffer, H.G. Benecke and W. Sender and his crew in the Drömling Nature Park for their essential help in the field work in Germany, H. Schmid, H. Eggers, G. Sterzer and N. Aljadeff for their help in the data downloading, O. Hatzofe from the Israel Nature and Parks Authority and B. Keeves from the Max Planck Institute of Animal Behavior for their help in retrieving lost transmitters and W. Heidrich and F. Kuemmeth from e-obs GmbH for their dedicated technical support. We would like to thank J. Shamoun-Baranes and an anonymous reviewer that substantially improved the manuscript with their comments and suggestions. We acknowledge the generous funding of DIP grants (DFG) NA 846-1-1 and WI 3576-1-1 to R.N., F.J. and M.W. This study was also supported by the Minerva Center for Movement Ecology granted to R.N. and the Max Planck Institute of Animal Behavior. S.R. was supported by a doctoral bird study scholarship of the Ministry of Science and Technology, Israel.

## Authors’ contributions

P.B. and N.S. conceived the study and designed the methodology with the help of S.R., N.H. and R.N. S.R. and M.K. carried out the field work with the help of M.W. and D.Z. P.B. analysed the data and compiled all figures. P.B. and N.S. led the writing of the manuscript and all authors contributed to the revisions of the draft. S.R., N.H. and R.N. further contributed to data interpretation and manuscript revision. All authors contributed critically to the drafts and gave final approval for publication.

## Data availability statement

The GPS-ACC data that was used in this study is available in Movebank Data Repository (movebank.org): https://doi.org/10.5441/001/1.v8d24552 (Rotics et al., 2018), and https://doi.org/10.5441/001/1.hn1bd23k (Rotics et al., 2016).

Rotics, S., Kaatz, M., Resheff, Y. S., Turjeman, S. F., Zurell, D., Sapir, N., … Nathan, R. (2016). Data from: The challenges of the first migration: movement and behavior of juvenile versus adult white storks with insights regarding juvenile mortality. *Movebank Data Repository*. doi: 10.5441/001/1.hn1bd23k.

Rotics, S., Kaatz, M., Turjeman, S., Zurell, D., Wikelski, M., Sapir, N., … Nathan, R. (2018). Data from: Early arrival at breeding grounds: causes, costs and a trade-off with overwintering latitude. *Movebank Data Repository*. doi: https://doi.org/10.5441/001/1.v8d24552

