## Supplementary material for "Causes and consequences of facultative sea crossing in a soaring migrant": Table S; Figure S

### Supporting Methods

#### Environmental data annotation

Each GPS position was annotated with environmental data using the Env-DATA track annotation tool of MoveBank (Dodge et al., 2013). These data included (1) E-W wind (also known as the U wind component, with positive values indicating eastward wind), (2) N-S wind (the V component of the wind, with positive values indicating northward wind) that were downloaded for both 10 m above the surface and at the height of the storks' flight with temporal resolution of 3 hours, (3) thermal uplift velocity (6 hours temporal resolution, see the formula in Bohrer et al., 2012), and (4) surface air temperature (6 hours temporal resolution). The source of these data is the European Centre for Medium-Range Weather Forecasts (ECMWF) re-analysis data archive with 0.75 degrees of spatial resolution. From the U and V components of the wind we calculated wind speed ($W_s$) and direction (blowing towards $\theta$) as follows: $W_s = \sqrt{U^2 + V^2}$, and $\theta = \tan^{-1}(U, V)$. For each track segment between two positions we additionally calculated wind support ($W_p$), which represents the component of the wind vector towards the direction of the bird (positive values) or against it (negative values), by applying the following formula: $W_p = W_s \times \cos(\theta - \sigma_i)$, where $\sigma$ is the angle of each segment $i$ per individual (see below). Also, we calculated the crosswind $W_c$, which is the lateral component of the wind relative to the birds' tracks for each segment: $W_c = W_s \times \sin(\theta - \sigma_i)$. We used the U and V wind components at 10 meters above the surface because they were highly correlated with those estimated at the storks' heights (Spearman-ρ > 0.9, $p$ < 0.001).

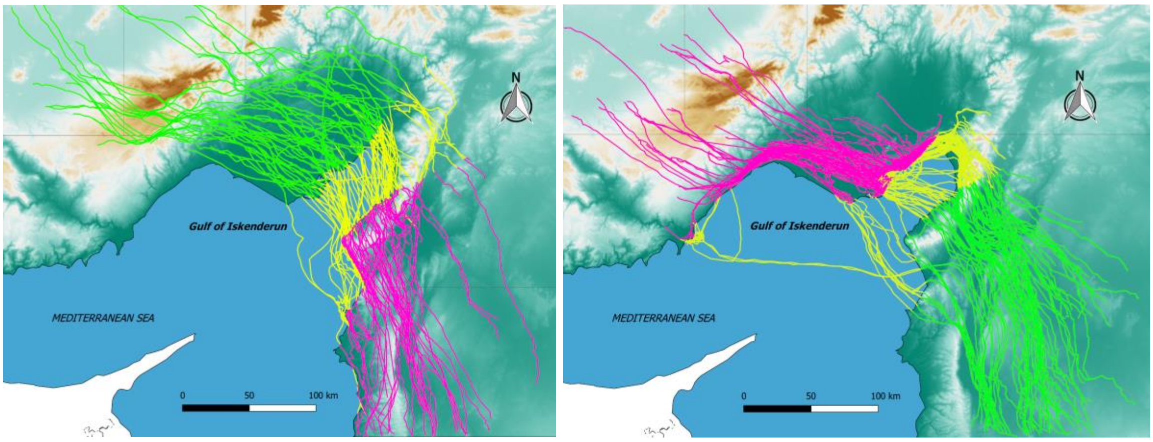


**Figure S1.** Illustration of the division of the storks' flight paths to 3 sections in relation to the bay: BEFORE (pink), ACROSS (yellow), and AFTER (green), in spring (left) and autumn (right).

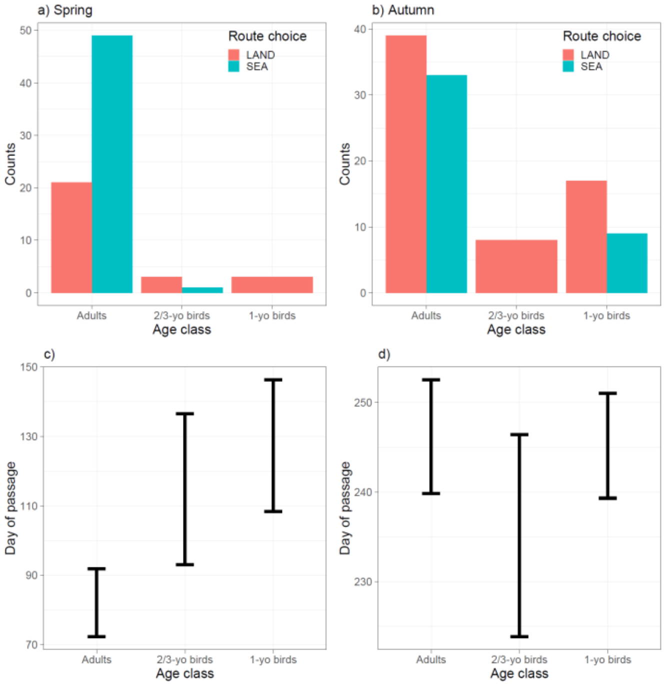


**Figure S2.** Frequencies and days of passage of three age classes of white storks that were analysed in the present study. The graphs show similar behaviour in sea crossing (a, b) and day of passage over the study area (c, d) between birds in their 1$^{st}$ year and birds in their 2$^{nd}$ and 3$^{rd}$ year (especially in spring), as well as differences between the behaviour of immature birds and adults.

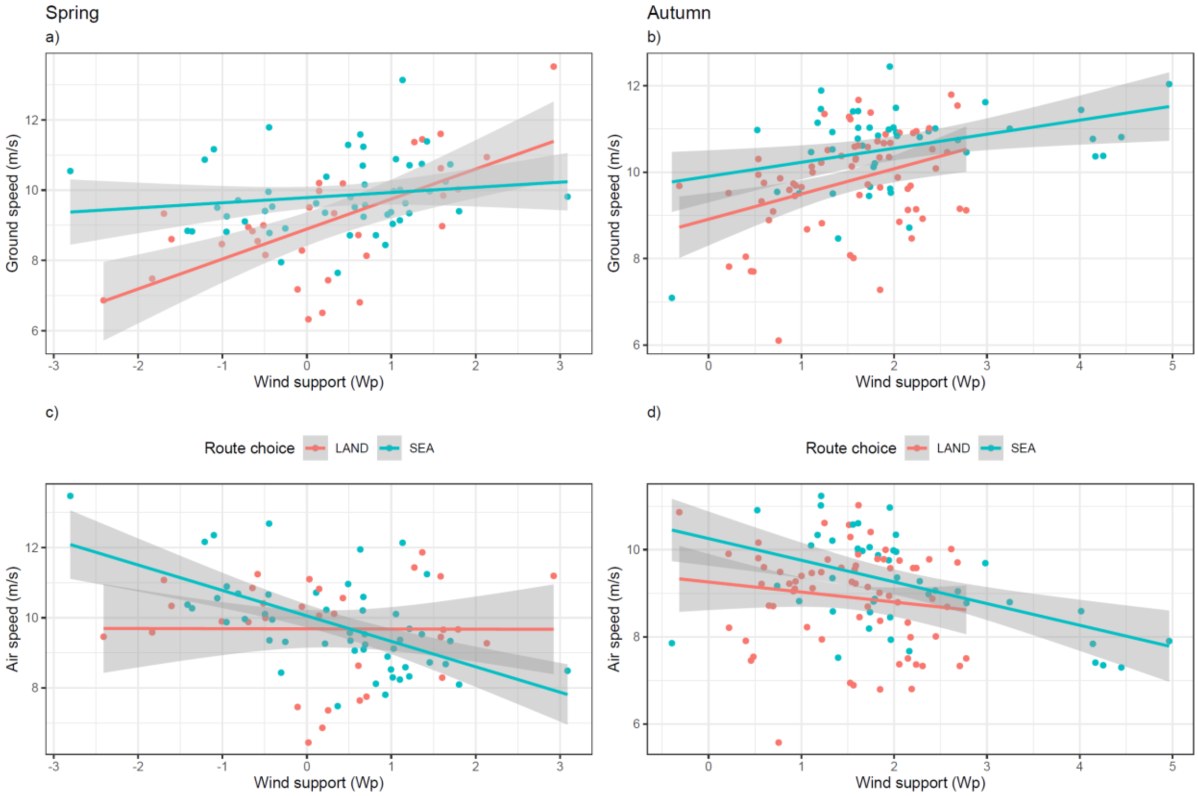


**Figure S3** – The effects of the interaction between route choice (LAND and SEA) and wind support (Wp) on ground (a, b) and air (c, d) speed in white storks during spring (a, c) and autumn (b, d) migration. Negative values are headwinds and positive values are tailwinds.

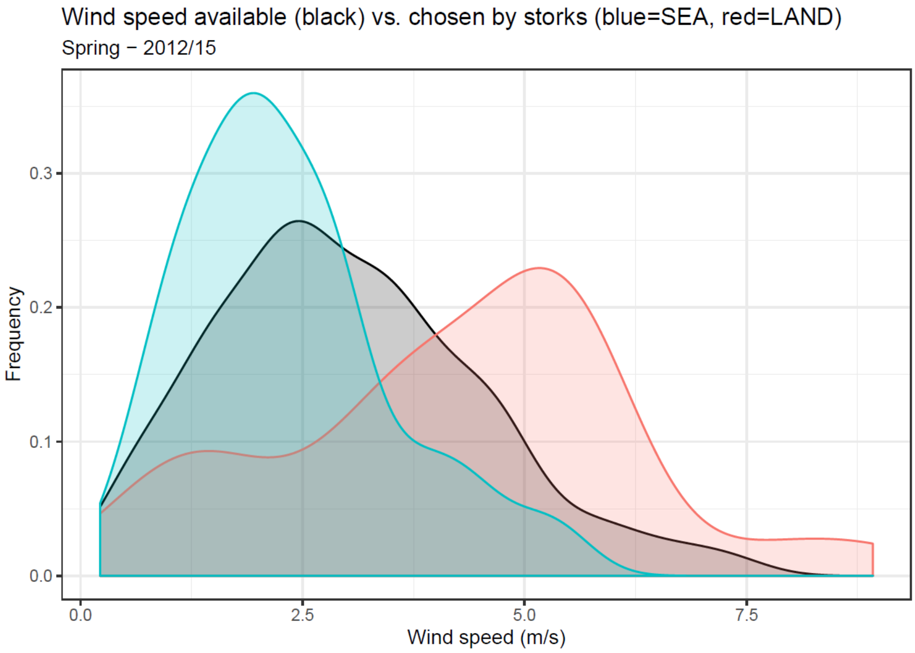


**Figure S4** – The wind speed available in the study area (black) during the entire spring migration period of white storks, and the speed of the wind used by the storks before crossing the bay (blue) and before undertaking a land detour (red).

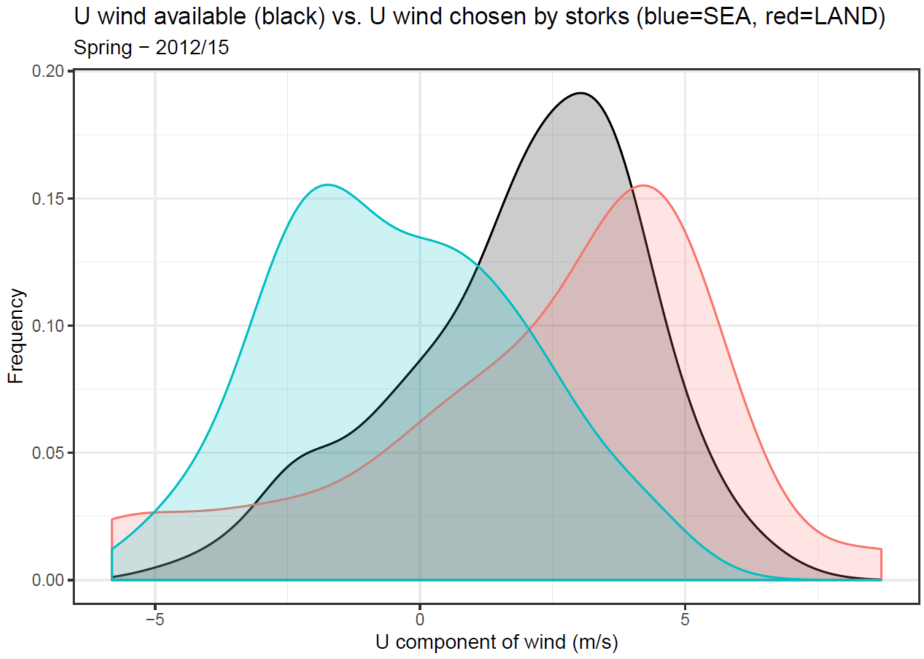


**Figure S5** – The U wind component speed available in the study area (black) during the entire spring migration period of white storks, and the U wind component speed used by the storks before crossing the bay (blue) and before undertaking a land detour (red). Positive values of the U wind component indicate eastward winds and negative ones indicate westward winds.

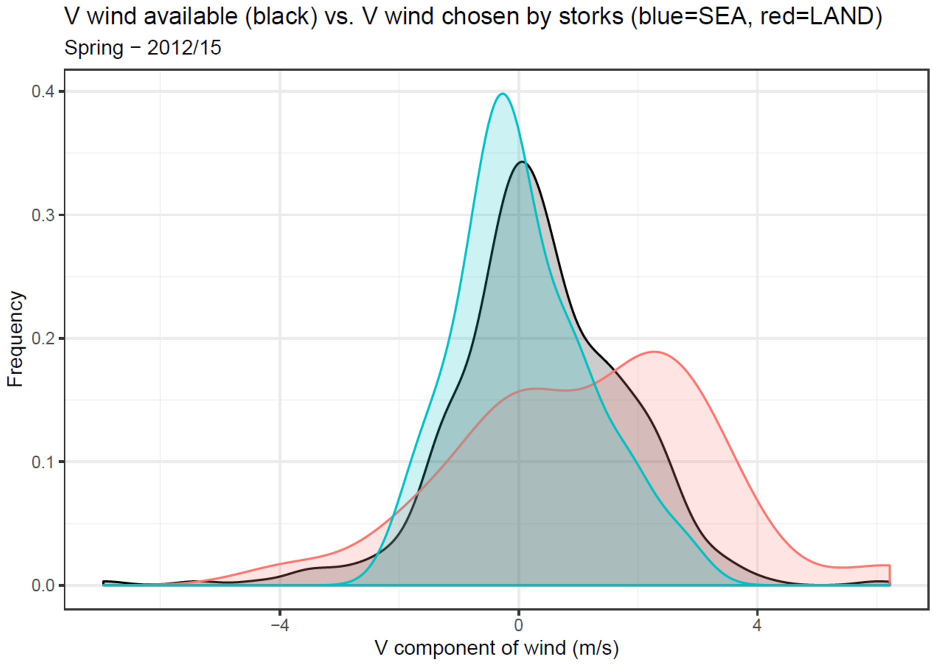


**Figure S6** – The V wind component speed available in the study area (black) during the entire spring migration period of white storks, and the V wind component speed used by the storks before crossing the bay (blue) or before undertaking a land detour (red). Positive values of the V component indicate northward winds and negative ones indicate southward winds.

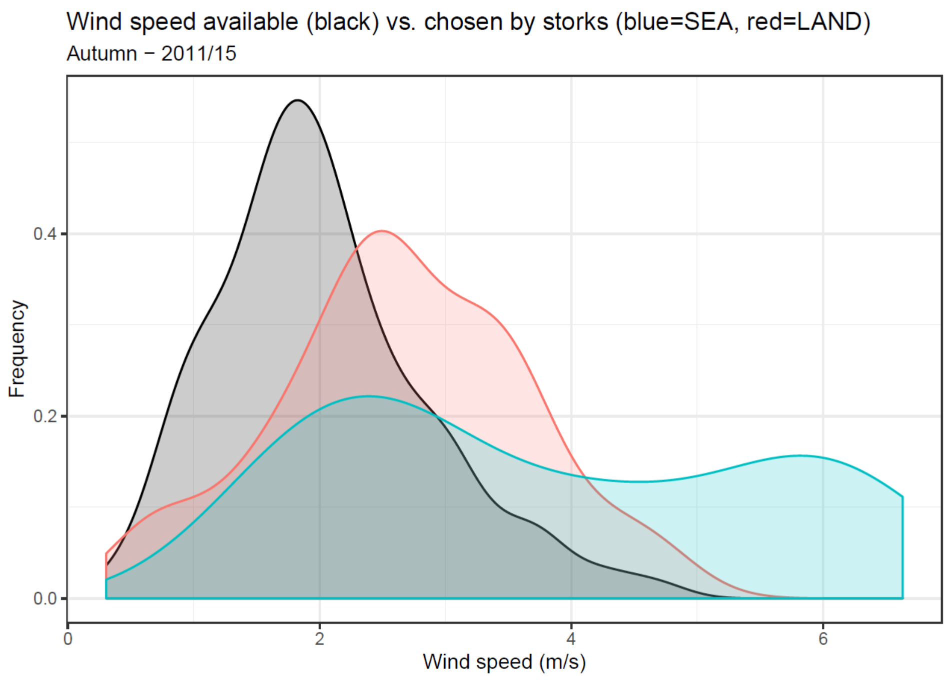


**Figure S7** – The wind speed available in the study area (black) during the entire autumn migration period of white storks, and the speed of the wind used by the storks before crossing the bay (blue) or before undertaking a land detour (red).

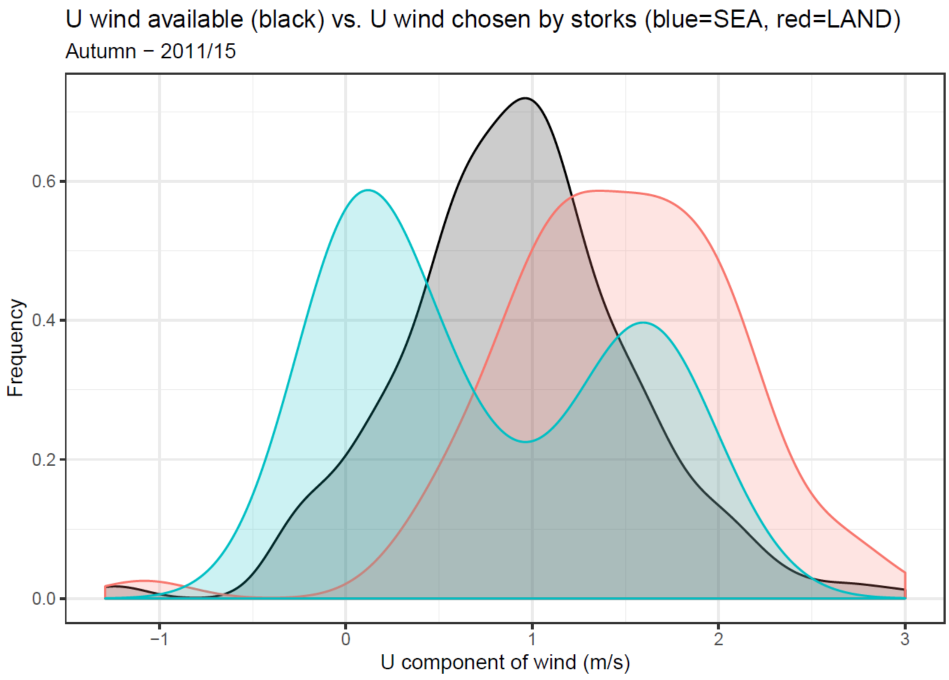


**Figure S8** – The U wind component speed available (black) during the entire autumn migration period of white storks, and the U wind component speed used by the storks before crossing the bay (blue) or before undertaking a land detour (red). Positive values of the U wind component indicate eastward winds and negative ones indicate westward winds.

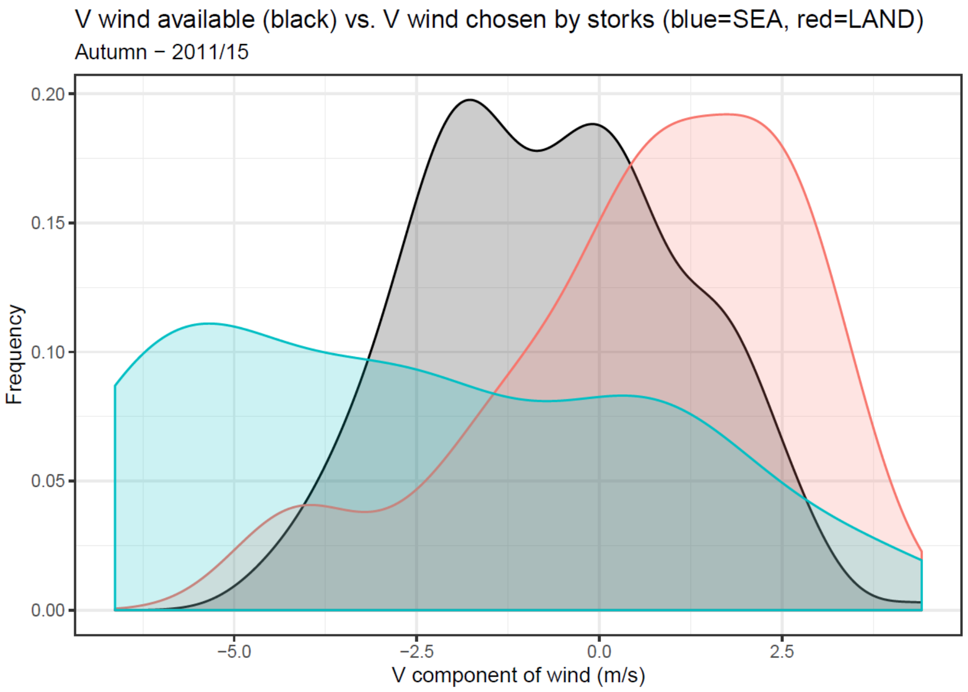


**Figure S9** – The V wind component speed available (black) during the entire autumn migration period of white storks, and the V wind component speed used by the storks before crossing the bay (blue) or before undertaking a land detour (red). Positive values of the V component indicate northward winds and negative ones indicate southward winds.

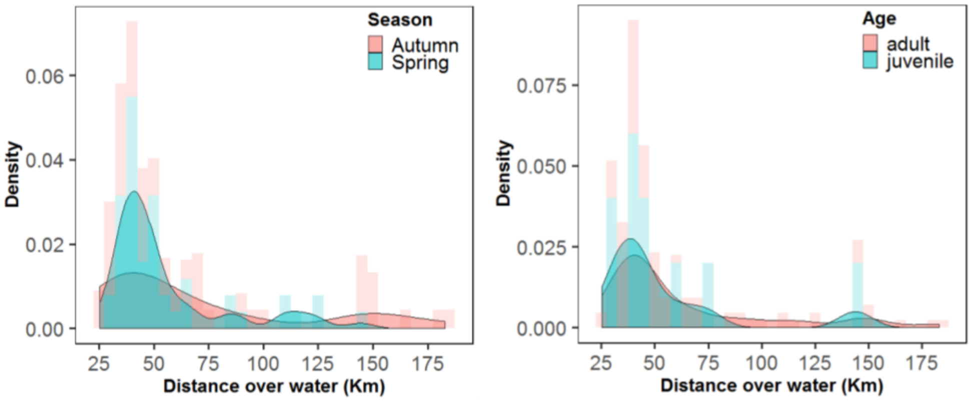


**Figure S10** – The distributions of the distances covered over the sea by white storks in relation to season (left) and age (right).

**Table S1** – Summary of pairwise comparisons of flight parameters between SEA and LAND groups of tracks in the 3 sections of their path over the study area relative to the bay. ODBA and proportion of active flight values are log-transformed to normalise their distribution.

| Season | Flight parameter | Path section relative to the bay | LAND - SEA Mean (s.d.) | β difference (s.e.) | *t* ratio | P value |
|---|---|---|---|---|---|---|
| Spring | Time flying (hours) | Before | 3.03 (1.78) - 3.16 (1.71) | -0.13 (0.42) | -0.31 | 0.75 |
| | | Across | 1.68 (0.98) - 1.31 (0.93) | 0.37 (0.41) | 0.91 | 0.36 |
| | | After | 2.71 (2.02) - 3.87 (2.5) | -1.16 (0.43) | -2.72 | **< 0.01** |
| | | Total | 6.80 (1.73) - 8.03 (2.03) | -1.26 (0.47) | -2.67 | **< 0.01** |
| | Distance covered (km) | Before | 93.4 (65.6) - 104.0 (60.8) | -10.1 (14.7) | -0.69 | 0.49 |
| | | Across | 57.8 (19.9) - 55.5 (27.6) | 2.36 (14.2) | 0.17 | 0.87 |

| | | | | | | |
|---|---|---|---|---|---|---|
| | | After | 91.1 (73.8) - 133 (87.9) | -42.1 (14.9) | -2.83 | **< 0.01** |
| | | Total | 222 (61.7) - 282 (67.7) | -56 (15.6) | -3.56 | **< 0.001** |
| | ODBA ($m^2 \cdot s^{-2}$) | Before | 2.88 (0.92) - 3.65 (1.47) | -0.18 (0.1) | -1.78 | 0.07 |
| | | Across | 3.71 (1.17) - 6.1 (1.84) | -0.52 (0.1) | -5.26 | **< 0.001** |
| | | After | 3.24 (2.44) - 3.28 (1.51) | -0.1 (0.1) | -0.96 | 0.33 |
| | | Total | 3.11 (0.93 - 3.8 (1.1) | -0.21 (0.07) | -3.12 | **< 0.01** |
| | Proportion of active flight | Before | 0.23 (0.14) - 0.36 (0.19) | -0.1 (0.05) | -2.14 | **< 0.05** |
| | | Across | 0.37 (0.23) - 0.75 (0.19) | -0.36 (0.05) | -7.64 | **< 0.001** |
| | | After | 0.35 (0.29) - 0.33 (0.17) | 0.03 (0.05) | 0.53 | 0.6 |
| | | Total | 0.29 (0.14) - 0.41 (0.14) | -0.11 (0.03) | -3.34 | **< 0.01** |
| | Ground speed ($m \cdot s^{-1}$) | Before | 7.94 (1.85) - 8.79 (1.69) | -0.75 (0.56) | -1.33 | 0.186 |
| | | Across | 10.9 (3.80) - 13 (2.75) | -1.92 (0.55) | -3.52 | **< 0.001** |
| | | After | 9.12 (2.75) - 9.37 (1.29) | -0.16 (0.57) | -0.28 | 0.782 |
| | | Total | 9.10 (1.65) - 9.84 (1.04) | -0.64 (0.31) | -2.08 | **< 0.05** |
| | Air speed ($m \cdot s^{-1}$) | Before | 8.41 (2.14 - 8.76 (2.01) | -0.24 (0.53) | -0.44 | 0.66 |
| | | Across | 11.0 (3.11) - 12.8 (2.45) | -1.65 (0.51) | -3.20 | **< 0.01** |
| | | After | 10.3 (2.17) - 9.11 (1.70) | 1.30 (0.54) | 2.40 | **< 0.05** |
| | | Total | 9.68 (1.44) - 9.80 (1.33) | -0.03 (0.32) | -0.08 | 0.93 |
| Autumn | Time flying (hours) | Before | 3.69 (2.24) - 2.95 (1.49) | 0.74 (0.3) | 2.50 | **< 0.05** |
| | | Across | 1.98 (0.53) - 1.70 (1.31) | 0.29 (0.3) | 0.95 | 0.34 |
| | | After | 3.61 (1.34) - 3.67 (1.19) | -0.06 (0.3) | -0.20 | 0.84 |
| | | Total | 8.26 (1.42) - 8.26 (1.68) | 0.05 (0.31) | 0.17 | 0.86 |
| | Distance covered (km) | Before | 123 (81.3) - 97.6 (53.6) | 25.47 (11.2) | 2.27 | **< 0.05** |
| | | Across | 59.8 (16.2) - 70.5 (47.5) | -10.73 (11.1) | -0.97 | 0.33 |
| | | After | 151 (58.6) - 148 (49.3) | 2.92 (11.4) | 0.25 | 0.80 |
| | | Total | 294 (62.6) - 314 (67.0) | -17.2 (13.4) | -1.29 | 0.20 |
| | ODBA ($m^2 \cdot s^{-2}$) | Before | 1.85 (0.5) - 2.52 (2.29) | -0.18 (0.1) | -1.78 | 0.08 |
| | | Across | 3.52 (1.28) - 6.32 (2.03) | -0.52 (0.1) | -5.26 | **< 0.001** |
| | | After | 2.05 (0.74) - 2.29 (1.69) | -0.1 (0.1) | -0.96 | 0.33 |
| | | Total | 2.28 (0.5) - 2.84 (0.8) | -0.21 (0.05) | -4.21 | **< 0.001** |
| | Proportion of active flight | Before | 0.1 (0.07) - 0.16 (0.22) | -0.07 (0.03) | -2.15 | **< 0.05** |
| | | Across | 0.32 (0.17) - 0.72 (0.2) | -0.4 (0.03) | -12.23 | **< 0.001** |
| | | After | 0.13 (0.12) - 0.15 (0.19) | -0.03 (0.03) | -1.03 | 0.3 |
| | | Total | 0.16 (0.08) - 0.24 (0.13) | -0.08 (0.02) | -3.78 | **< 0.001** |
| | Ground speed ($m \cdot s^{-1}$) | Before | 8.87 (1.21) - 9.10 (1.64) | -0.23 (0.29) | -0.79 | 0.43 |
| | | Across | 8.52 (1.32) - 12.3 (1.54) | -3.77 (0.28) | -13.27 | **< 0.001** |
| | | After | 11.4 (1.47) - 11.2 (1.41) | 0.28 (0.29) | 0.95 | 0.34 |
| | | Total | 9.79 (1.13) - 10.6 (0.98) | -0.77 (0.23) | -3.32 | **< 0.01** |
| | Air speed ($m \cdot s^{-1}$) | Before | 8.06 (1.39) - 8.65 (2.41) | -0.59 (0.35) | -1.69 | 0.09 |
| | | Across | 7.9 (1.53) - 10.4 (1.69) | -2.51 (0.35) | -7.26 | **< 0.001** |
| | | After | 10.3 (1.67) - 9.24 (1.67) | 1.01 (0.36) | 2.83 | **< 0.01** |
| | | Total | 8.91 (1.13) - 9.22 (1.09) | -0.3 (0.24) | -1.25 | 0.22 |

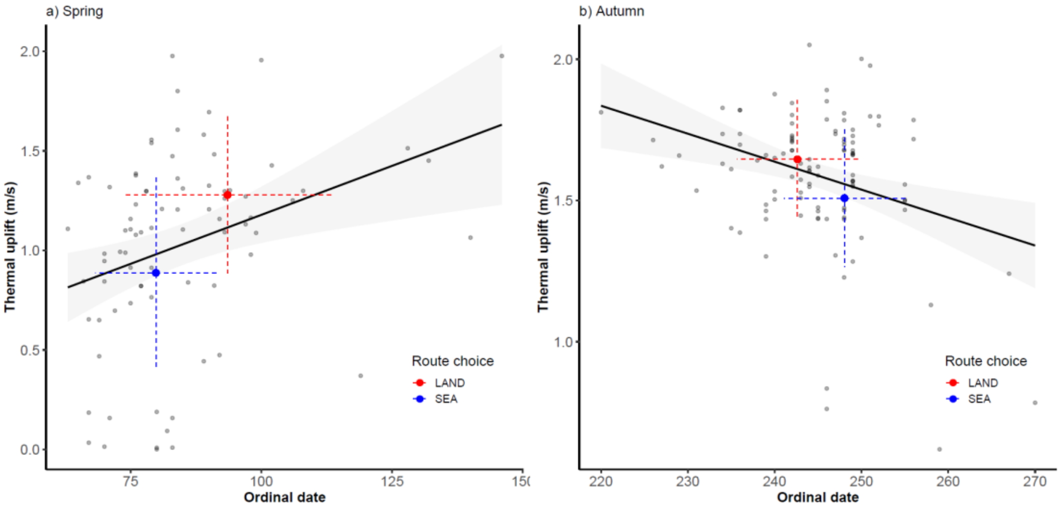


**Figure S11** – The relationships between thermal uplift and ordinal date (number of days from the 1st January). Regression line is in black. Blue and red dots are mean values for the sea-crossing and land detour choices, respectively. Dashed lines are standard deviations.

### Route choice analysis

#### Spring – GLMM

Full model formula: Route_choice ~ Temperature + Wind_speed + EW_wind + Ordinal_date + Sex + Age + (1|Bird_ID) + (1|Year)

Number of obs. = 77

**Table S2** – Variables selected for the full model after checking for autocorrelation and collinearity of the most biologically meaningful variables.

| Model structure | Variable name | Variable type | N. levels (range or values) | Unit system |
|---|---|---|---|---|
| Dependent | Route choice | Binomial | 2 (0, 1) | – |
| Explanatory | Wind speed (*Ws*) | Continuous | – | $m \cdot s^{-1}$ |
| Explanatory | N-S wind | Continuous | – | $m \cdot s^{-1}$ |
| Explanatory | E-W wind | Continuous | – | $m \cdot s^{-1}$ |
| Explanatory | Ordinal date | Continuous | – | – |
| Explanatory | Age | Categorical | 2 (adult, 1to3-yo*) | – |
| Explanatory | Sex | Categorical | 2 (female, male) | – |
| Random | Bird ID | Factor | 41 | – |
| Random | Year | Factor | 4 (2012-2015) | – |

* 1to3-yo = birds from 1 to 3 years old.

**Table S3** – Selected (ΔAICc < 7) generalized linear mixed models of environmental variables affecting the route choice in migrating White storks. First 10 models out of 23 are shown.

| Model | Variables | *w* | AICc | ΔAICc |
|---|---|---|---|---|
| 1 | Wind speed, E-W wind, Ordinal date | 0.21 | 62.25 | 0.00 |
| 2 | Wind speed, E-W wind, Ordinal date, Age | 0.15 | 62.87 | 0.62 |
| 3 | Wind speed, E-W wind, Ordinal date, Temperature | 0.07 | 64.37 | 2.12 |
| 4 | Wind speed, E-W wind, Ordinal date, N-S wind | 0.07 | 64.55 | 2.30 |
| 5 | Wind speed, E-W wind, Sex | 0.06 | 64.65 | 2.40 |
| 6 | Wind speed, E-W wind, Age | 0.06 | 64.73 | 2.48 |
| 7 | Wind speed, E-W wind, Ordinal date, Age, Temperature | 0.05 | 64.93 | 2.68 |
| 8 | Wind speed, E-W wind, Ordinal date, Age, Sex | 0.05 | 65.14 | 2.90 |
| 9 | Wind speed, E-W wind, Ordinal date, N-S wind, Age | 0.04 | 65.31 | 3.06 |
| 10 | Wind speed, E-W wind, Age, Sex | 0.04 | 65.73 | 3.48 |
| ... | ... | ... | ... | ... |

AICc: Akaike's information criterion corrected for sample size, ΔAICc: difference in AICc between a given model and the best model, *w*: Akaike's weights.

**Table S4** – Model-averaged (subset models ΔAICc < 7) coefficients (*β*) with 95% confidence intervals (LCI, UCI) of generalized linear mixed models of environmental variables affecting the route choice in migrating White storks, ranked by their predictive importance (Σ*w*). *N* indicates the number of models containing a given variable.

| Variable | *β* | LCI | UCI | *p* | Σ*w* | *N* |
|---|---|---|---|---|---|---|
| (Intercept) | 1.07 | 0.1 | 2.13 | 0.047 | – | – |
| Wind speed | –1.57 | –2.61 | –0.53 | **< 0.01** | 1 | 23 |
| E-W wind | –1.08 | –2.05 | –0.12 | **0.028** | 0.98 | 21 |
| Ordinal date | –1.35 | –2.48 | –0.23 | **0.018** | 0.85 | 17 |
| Age | –2.71 | –6.72 | 1.30 | 0.19 | 0.50 | 13 |
| Sex | 0.32 | –1.53 | 2.16 | 0.74 | 0.25 | 10 |
| Temperature | 0.27 | –0.78 | 1.31 | 0.62 | 0.24 | 9 |
| N-S wind | –0.14 | –0.97 | 0.69 | 0.74 | 0.23 | 10 |

**Table S5** – Best model: Route_choice ~ Wind_speed + EW_wind + Ordinal_date + (1 | Bird_ID) + (1|Year) and information about the properties of its factors.

| Variable | *β* | SE | *z* | *p* |
|---|---|---|---|---|
| (Intercept) | 0.97 | 0.38 | 2.57 | 0.01 |
| Wind speed | –1.55 | 0.46 | –3.35 | **< 0.001** |
| E-W wind | –1.11 | 0.42 | –2.64 | **< 0.01** |
| Ordinal date | –1.49 | 0.45 | –3.27 | **< 0.01** |

Random effects variance: Bird ID < 0.001, Year < 0.001

The figures (below) show the major effects on route choice as reported by the best model:

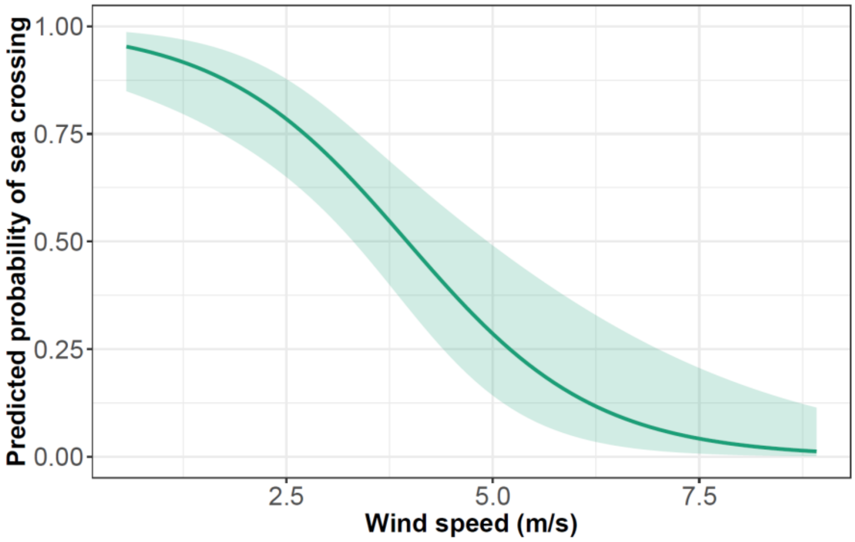


**Figure S12** – Logistic regression line with 95% CI (shaded area) of predicted probabilities of sea crossing in spring in relation to wind speed (m/s) in the section BEFORE the bay (see Table S5).

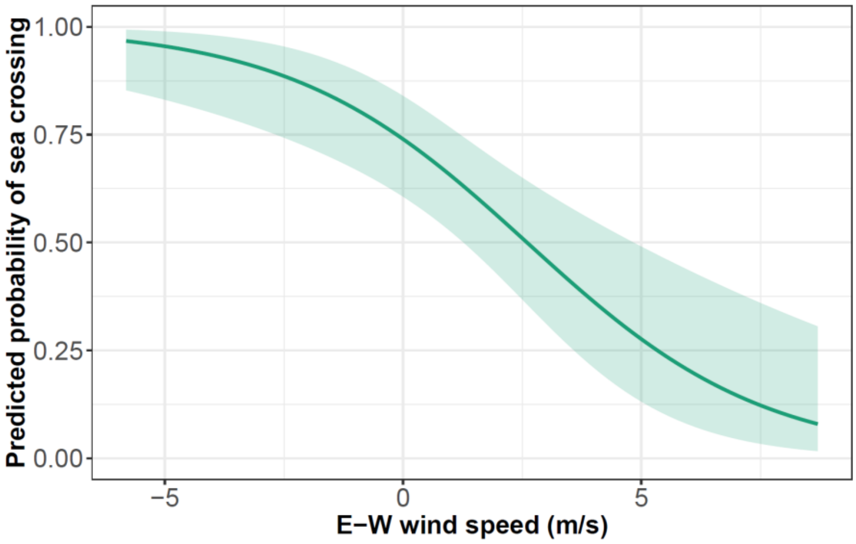


**Figure S13** – Logistic regression line with 95% CI (shaded area) of predicted probabilities of sea crossing in spring in relation to E-W wind component speed (m/s) in the section BEFORE the bay (see Table S5).

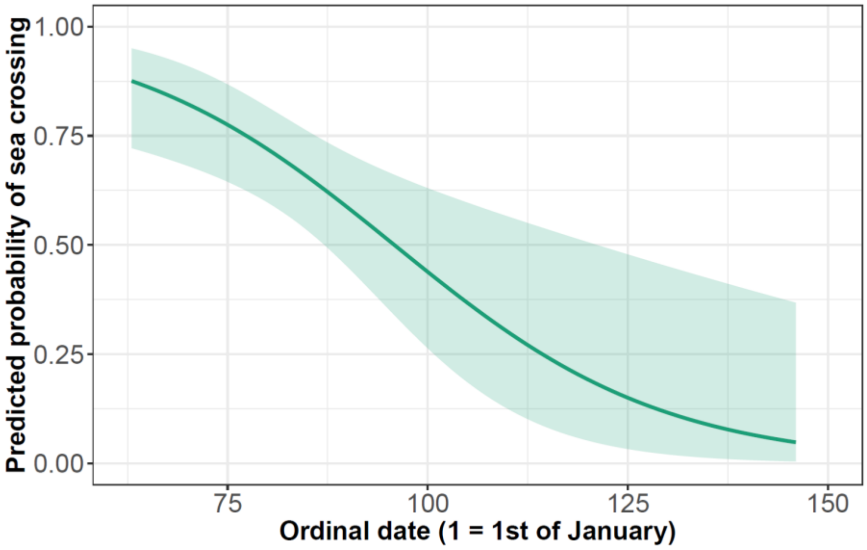


**Figure S14** – Logistic regression line with 95% CI (shaded area) of predicted probabilities of sea crossing in spring in relation to ordinal date (the day of stork passage over the bay area, counted from January 1st) in the section BEFORE the bay (see Table S5).

### Autumn – GLMM

Full model formula: Route_choice ~ Wind_speed + NS_wind + EW_wind + Thermal_uplift + Distance_from_bay + Ordinal_date + Sex + Age + (1|Bird_ID) + (1|Year)

Number of obs. = 106

**Table S6** – Variables selected for the full model after checking for autocorrelation and collinearity of the most biologically meaningful variables.

| **Model structure** | **Variable name** | **Variable type** | **N. levels (range or values)** | **Unit system** |
|---|---|---|---|---|
| Dependent | Route choice | Binomial | 2 (0, 1) | – |
| Explanatory | Wind speed (*Ws*) | Continuous | – | $m \cdot s^{-1}$ |
| Explanatory | N-S wind | Continuous | – | $m \cdot s^{-1}$ |
| Explanatory | E-W wind | Continuous | – | $m \cdot s^{-1}$ |
| Explanatory | Thermal uplift | Continuous | – | $m \cdot s^{-1}$ |
| Explanatory | Distance before bay | Continuous | – | km |
| Explanatory | Ordinal date | Continuous | – | – |
| Explanatory | Age | Categorical | 2 (adult, 1to3-yo*) | – |
| Explanatory | Sex | Categorical | 2 (female, male) | – |
| Random | Bird ID | Factor | 67 | – |
| Random | Year | Factor | 5 (2011-2015) | – |

* 1to3-yo = birds from 1 to 3 years old.

**Table S7** – Selected (ΔAICc < 7) generalized linear mixed models of environmental variables affecting the route choice in migrating White storks. First 10 models out of 69 are shown.

| Model | Variables | *w* | AICc | ΔAICc |
|---|---|---|---|---|
| 1 | Ordinal date, N-S wind, Age | 0.09 | 111.46 | 0.00 |
| 2 | Ordinal date, N-S wind | 0.08 | 111.73 | 0.26 |
| 3 | Ordinal date, N-S wind, Age, E-W wind | 0.05 | 112.91 | 1.44 |
| 4 | Ordinal date, N-S wind, Wind speed | 0.04 | 113.12 | 1.65 |
| 5 | Ordinal date, N-S wind, Age, Wind speed | 0.04 | 113.34 | 1.88 |
| 6 | Ordinal date, N-S wind, Distance covered before bay | 0.04 | 113.39 | 1.92 |
| 7 | Ordinal date, N-S wind, Age, Distance covered before bay | 0.04 | 113.42 | 1.96 |
| 8 | Ordinal date, N-S wind, E-W wind | 0.03 | 113.45 | 1.98 |
| 9 | Ordinal date, N-S wind, Age, Thermal uplift | 0.03 | 113.62 | 2.15 |
| 10 | Ordinal date, N-S wind, Age, Sex | 0.03 | 113.76 | 2.29 |
| ... | ... | ... | ... | ... |

AICc: Akaike's information criterion corrected for sample size, ΔAICc: difference in AICc between a given model and the best model, *w*: Akaike's weights.

**Table S8** – Model-averaged (subset models ΔAICc < 7) coefficients ($\beta$) with 95% confidence intervals (LCI, UCI) of generalized linear mixed models of environmental variables affecting the route choice in migrating White storks, ranked by their predictive importance ($\Sigma w$). *N* indicates the number of models containing the given variable.

| Variable | *β* | LCI | UCI | *p* | Σ*w* | *N* |
|---|---|---|---|---|---|---|
| (Intercept) | –0.38 | –1.05 | 0.29 | 0.27 | – | – |
| Ordinal date | 0.98 | 0.19 | 1.79 | **0.014** | 0.99 | 67 |
| N-S wind | –1.14 | –1.97 | –0.32 | < **0.01** | 0.94 | 56 |
| Age | –0.89 | –2.13 | 0.34 | 0.16 | 0.50 | 32 |
| Wind speed | 0.48 | –0.44 | 1.39 | 0.31 | 0.37 | 39 |
| E-W wind | –0.38 | –1.15 | 0.39 | 0.34 | 0.35 | 36 |
| Thermal uplift | –0.25 | –1.06 | 0.55 | 0.54 | 0.29 | 34 |
| Distance covered before bay | 0.17 | –0.43 | 0.76 | 0.58 | 0.27 | 28 |
| Sex | –0.04 | –1.10 | 1.03 | 0.94 | 0.22 | 26 |

**Table S9** – Best model: Route_choice ~ Ordinal date + NS_wind + (1 | bird_id) + (1 | year) and information about the properties of its factors.

| Variable | *β* | SE | *z* | *p* |
|---|---|---|---|---|
| (Intercept) | –0.28 | 0.29 | –0.93 | 0.35 |
| Ordinal date | 1.04 | 0.39 | 2.66 | < **0.01** |
| N-S wind | –1.27 | 0.36 | –3.55 | < **0.001** |

| | | | | |
|---|---|---|---|---|
| Age | –0.89 | 0.61 | –1.48 | 0.14 |

Random effects variance: Bird ID = 0.09, Year < 0.001

Figures (below) show the major effects on route choice as reported by the best model:

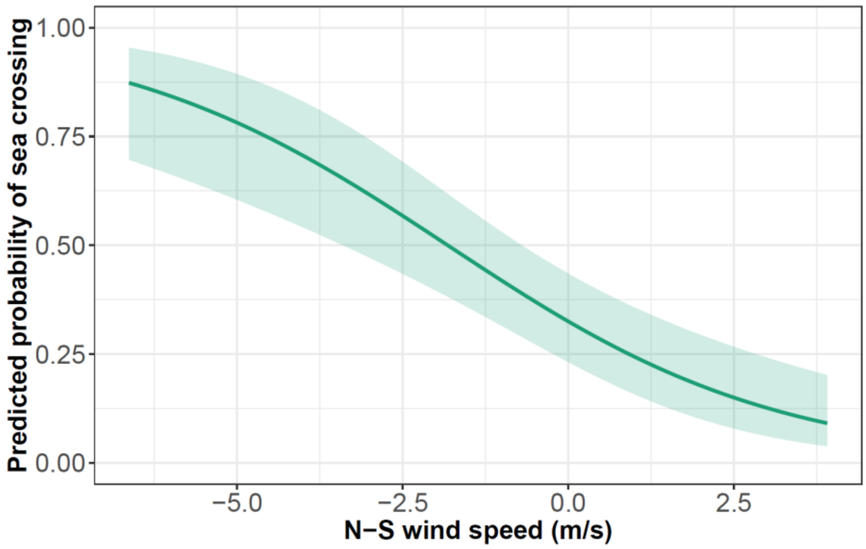


**Figure S15** – Logistic regression line with 95% CI (shaded area) of probabilities of sea crossing in autumn in relation to N-S wind component speed (m/s) in the section BEFORE the bay (see Table S9).

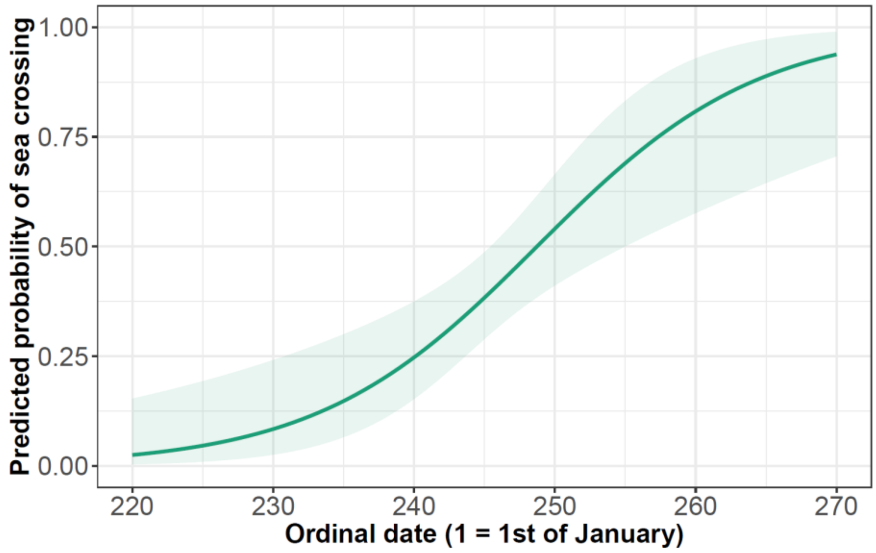


**Figure S16** – Logistic regression line with 95% CI (shaded area) of probabilities of sea crossing in autumn in relation to ordinal date (day of stork passage over the bay area) in the section BEFORE the bay (see Table S9).

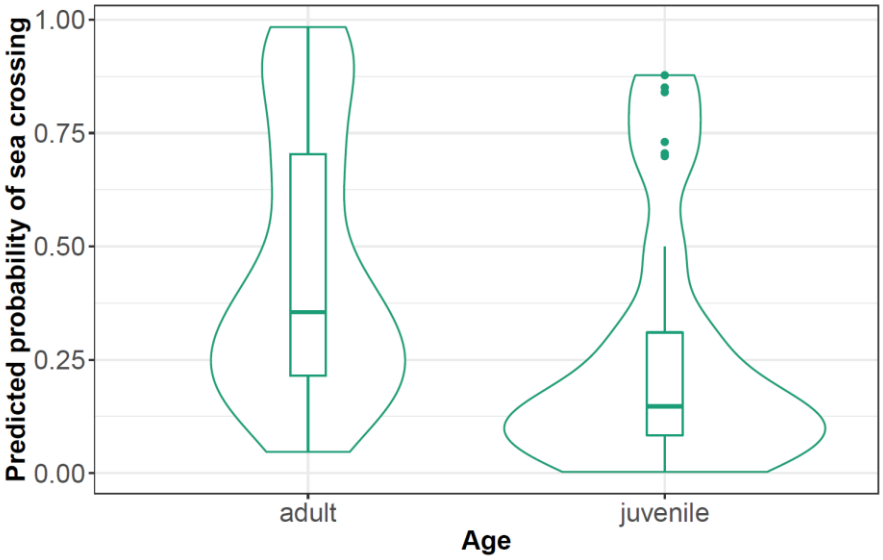


**Figure S17** – Sea-crossing probabilities in autumn in the two different age groups (see Table S9). Boxplots represent the median (horizontal line), first and third quartile (the upper and lower limits of the box), the range of the data (vertical line), outliers (points) and the data distribution (area around the boxplots).

### Flight speed analysis

#### Ground speed – Spring – LMM

Full model formula: Ground_speed ~ Route_choice × (Temperature + Wind_support + Crosswinds + Wind_speed + Thermal_uplift + Age + Sex) + (1|Bird_ID) + (1|Year)

Number of obs. = 83

**Table S10** – Variables selected for the full model after checking for autocorrelation and collinearity of the most biologically meaningful variables.

| **Model structure** | **Variable name** | **Variable type** | **N. levels (range or values)** | **Unit system** |
|---|---|---|---|---|
| Dependent | Ground speed | Continuous | – | $m{\cdot}s^{-1}$ |
| Explanatory | Route choice | Binomial | 2 (0, 1) | – |
| Explanatory | Temperature | Continuous | – | °C |
| Explanatory | Wind support (*Wp*) | Continuous | – | $m{\cdot}s^{-1}$ |
| Explanatory | Crosswinds (*Wc*) | Continuous | – | $m{\cdot}s^{-1}$ |
| Explanatory | Wind speed (*Ws*) | Continuous | – | $m{\cdot}s^{-1}$ |
| Explanatory | Thermal uplift | Continuous | – | $m{\cdot}s^{-1}$ |
| Explanatory | Sex | Categorical | 2 (female, male) | – |
| Explanatory | Age | Categorical | 2 (adult, 1to3-yo*) | – |
| Random | Bird ID | Factor | 35 | – |
| Random | Year | Factor | 4 | – |

* 1to3-yo = birds from 1 to 3 years old.

**Table S11** – Selected (ΔAICc < 7) generalized linear mixed models of environmental variables affecting the route choice in migrating White storks. First 10 models out of 59 are shown.

| Model | Variables | *w* | AICc | ΔAICc |
|---|---|---|---|---|
| 1 | Route choice × Wind support | 0.12 | 270.70 | 0.00 |
| 2 | Route choice × Wind support, Wind speed | 0.08 | 271.43 | 0.73 |
| 3 | Route choice × Wind support, Route choice × Age | 0.07 | 271.74 | 1.04 |
| 4 | Route choice × Wind support, Wind speed, Route choice × Age | 0.05 | 272.31 | 1.61 |
| 5 | Route choice × Wind support, Thermal uplift | 0.04 | 272.91 | 2.21 |
| 6 | Route choice × Wind support, Age | 0.04 | 273.03 | 2.33 |
| 7 | Route choice × Wind support, Wind speed, Age | 0.04 | 273.06 | 2.36 |
| 8 | Route choice × Wind support, Route choice × Sex | 0.03 | 273.15 | 2.45 |
| 9 | Route choice × Wind support, Wind speed, Thermal uplift | 0.03 | 273.23 | 2.53 |
| 10 | Route choice × Wind support, Route choice × Sex, Wind speed | 0.03 | 274.27 | 2.58 |
| ... | ... | ... | ... | ... |

AICc: Akaike's information criterion corrected for sample size, ΔAICc: difference in AICc between a given model and the best model, *w*: Akaike's weights.

**Table S12** – Model-averaged (subset models ΔAICc < 7) coefficients (*β*) with 95% confidence intervals (LCI, UCI) of generalized linear mixed models of environmental variables affecting the route choice in migrating White storks, ranked by their predictive importance (Σ*w*). *N* indicates the number of models containing the given variable.

| Variable | *β* | LCI | UCI | *p* | Σ*w* | *N* |
|---|---|---|---|---|---|---|
| (Intercept) | 9.01 | 8.29 | 9.72 | < 0.001 | – | – |
| Route choice | 0.79 | –0.00 | 1.58 | 0.051 | 1 | 59 |
| Wind support | 0.92 | 0.51 | 1.32 | **< 0.001** | 1 | 59 |
| Route choice × Wind support | –0.74 | –1.24 | –0.24 | **0.003** | 0.96 | 40 |
| Wind speed | 0.31 | –0.01 | 0.63 | 0.057 | 0.40 | 25 |
| Age | 0.02 | –1.00 | 1.05 | 0.965 | 0.35 | 21 |
| Thermal uplift | 0.28 | –0.07 | 0.63 | 0.114 | 0.25 | 21 |
| Sex | 0.45 | –0.39 | 1.29 | 0.293 | 0.23 | 19 |
| Route choice × Age | 1.44 | –0.98 | 3.85 | 0.244 | 0. 18 | 6 |
| Route choice × Sex | –0.89 | –1.89 | 0.12 | 0.083 | 0.12 | 8 |
| Crosswind | 0.15 | –0.14 | 0.44 | 0.301 | 0.09 | 10 |
| Temperature | 0.07 | –0.30 | 0.45 | 0.709 | 0.08 | 12 |
| Route choice × Thermal uplift | –0.40 | –0.96 | 0.15 | 0.149 | 0.05 | 4 |
| Route choice × Wind speed | –0.17 | –0.77 | 0.43 | 0.583 | 0.04 | 5 |

| | | | | | | |
|---|---|---|---|---|---|---|
| Route choice × Temperature | 0.39 | –0.15 | 0.94 | 0.156 | 0.02 | 3 |
| Route choice × Crosswind | 0.17 | –0.38 | 0.71 | 0.549 | 0.01 | 1 |

**Table S13** – Best model: Groundspeed ~ Route choice × Wind support + (1 | Bird_ID) + (1 | year) and information about the properties of its factors.

| Variable | *β* | SE | t-value | *p* |
|---|---|---|---|---|
| (Intercept) | 9.19 | 0.28 | 32.42 | < 0.001 |
| Route choice | 0.54 | 0.25 | 2.13 | **0.035** |
| Wind support | 0.98 | 0.18 | 5.41 | **< 0.001** |
| Route choice × Wind support | –0.78 | 0.24 | –3.18 | **0.002** |

Random effects variance: Bird ID < 0.001, Year = 0.156

### Ground speed – Autumn – LMM

Full model formula: Ground_speed ~ Route_choice × (Temperature + Wind_support + Crosswind + Thermal_uplift + Ordinal_date + Sex + Age) + (1|Bird_ID) + (1|Year)

Number of obs. = 113

**Table S14** – Variables selected for the full model after checking for autocorrelation and collinearity of the most biologically meaningful variables.

| Model structure | Variable name | Variable type | N. levels (range or values) | Unit system |
|---|---|---|---|---|
| Dependent | Ground speed | Continuous | _ | $m \cdot s^{-1}$ |
| Explanatory | Route choice | Binomial | 2 (0, 1) | – |
| Explanatory | Temperature | Continuous | – | °C |
| Explanatory | Wind support (*Wp*) | Continuous | – | $m \cdot s^{-1}$ |
| Explanatory | Crosswind (*Wc*) | Continuous | – | $m \cdot s^{-1}$ |
| Explanatory | Wind speed (*Ws*) | Continuous | – | $m \cdot s^{-1}$ |
| Explanatory | Thermal uplift | Continuous | – | $m \cdot s^{-1}$ |
| Explanatory | Ordinal date | Continuous | – | – |
| Explanatory | Age | Categorical | 2 (adult, 1to3-yo*) | – |
| Explanatory | Sex | Categorical | 2 (female, male) | – |
| Random | Bird ID | Factor | 67 | – |
| Random | Year | Factor | 5 | – |

* 1to3-yo = birds from 1 to 3 years old.

**Table S15** – Selected (ΔAIC < 7) generalized linear mixed models of environmental variables affecting the route choice in migrating White storks. First 10 models out of 73 are shown.

| Model | Variables | *w* | AICc | ΔAICc |
|---|---|---|---|---|
| 1 | Route choice, Thermal uplift, Wind speed, Age | 0.09 | 313.39 | 0.00 |
| 2 | Route choice × Wind speed, Thermal uplift, Age | 0.09 | 313.51 | 0.12 |
| 3 | Route choice, Thermal uplift, Wind speed, Age, Crosswind | 0.08 | 313.61 | 0.22 |
| 4 | Route choice × Wind speed, Thermal uplift, Route choice × Age | 0.05 | 314.71 | 1.32 |
| 5 | Route choice, Wind speed, Age, Crosswind | 0.04 | 315.14 | 1.75 |
| 6 | Route choice × Thermal uplift, Wind speed, Age | 0.04 | 315.15 | 1.76 |
| 7 | Route choice × Age, Thermal uplift, Wind speed | 0.04 | 315.26 | 1.87 |
| 8 | Route choice, Thermal uplift, Crosswind, Wind speed | 0.03 | 315.31 | 1.92 |
| 9 | Route choice, Crosswind, Wind speed | 0.03 | 315.66 | 2.27 |
| 10 | Route choice × Age, Thermal uplift, Crosswind, Wind speed | 0.03 | 315.71 | 2.32 |
| ... | ... | ... | ... | ... |

AIC: Akaike's information criterion, ΔAIC: difference in AIC between a given model and the best model, *w*: Akaike's weights.

**Table S16** – Model-averaged (subset models ΔAICc < 7) coefficients (*β*) with 95% confidence intervals (LCI, UCI) of generalized linear mixed models of environmental variables affecting the route choice in migrating White storks, ranked by their predictive importance (Σ*w*). *N* indicates the number of models containing the given variable.

| Variable | *β* | LCI | UCI | *p* | Σ*w* | *N* |
|---|---|---|---|---|---|---|
| (Intercept) | 9.92 | 9.55 | 10.30 | < 0.001 | – | – |
| Route choice | 0.78 | 0.30 | 1.26 | **< 0.01** | 1 | 73 |
| Wind speed | 0.43 | 0.18 | 0.68 | **< 0.001** | 0.99 | 71 |
| Thermal uplift | 0.36 | 0.10 | 0.63 | **0.007** | 0.85 | 56 |
| Age | –0.46 | –0.86 | –0.06 | **0.025** | 0.79 | 49 |
| Crosswind | –0.27 | –0.53 | –0.02 | **0.037** | 0.45 | 36 |
| Route choice × Wind speed | –0.31 | –0.73 | 0.09 | 0.13 | 0.24 | 14 |
| Route choice × Age | –0.33 | –1.12 | 0.47 | 0.42 | 0.16 | 11 |
| Route choice × Thermal uplift | –0.23 | –0.61 | 0.14 | 0.22 | 0.13 | 10 |
| Sex | –0.18 | –0.62 | 0.27 | 0.44 | 0.12 | 16 |
| Ordinal date | 0.12 | –0.07 | 0.31 | 0.23 | 0.07 | 10 |
| Route choice × Sex | 0.65 | –0.03 | 1.33 | 0.06 | 0.05 | 6 |
| Wind support | 0.06 | –0.21 | 0.33 | 0.66 | 0.05 | 9 |
| Route choice × Crosswind | 0.20 | –0.38 | 0.79 | 0.49 | 0.05 | 6 |
| Temperature | –0.06 | –0.30 | 0.17 | 0.61 | 0.05 | 9 |
| Route choice × Wind support | –0.33 | –0.68 | 0.01 | 0.06 | 0.01 | 1 |
| Route choice × Temperature | 0.27 | –0.10 | 0.65 | 0.15 | 0.01 | 1 |
| Route choice × Ordinal date | –0.17 | –0.54 | 0.21 | 0.38 | 0.01 | 1 |

**Table S17** – Best model: Groundspeed ~ Route choice + Crosswind + Thermal uplift + Age + (1 | bird_id) + (1 | year) and information about the properties of its factors.

| Variable | *β* | SE | t-value | *p* |
|---|---|---|---|---|
| (Intercept) | 9.96 | 0.15 | 66.47 | < 0.001 |
| Route choice | 0.74 | 0.20 | 3.62 | **< 0.001** |
| Wind speed | 0.36 | 0.09 | 3.73 | **< 0.001** |
| Thermal uplift | 0.37 | 0.10 | 3.61 | **< 0.001** |
| Age | –0.49 | 0.19 | –2.60 | **0.011** |

Random effects variance: Bird ID = 0.06, Year = 0.013

### Air speed – Spring – LMM

Full model formula: Air_speed ~ Route_choice × (Temperature + Wind_speed + Wind_support + Crosswind + Thermal_uplift + Sex + Age) + (1|Bird_ID) + (1|Year)

Number of obs. = 83

**Table S18** – Variables selected for the full model after checking for autocorrelation and collinearity of the most biologically meaningful variables.

| Model structure | Variable name | Variable type | N. levels (range or values) | Unit system |
|---|---|---|---|---|
| Dependent | Air speed | Continuous | – | $m \cdot s^{-1}$ |
| Explanatory | Route choice | Binomial | 2 (0, 1) | – |
| Explanatory | Temperature | Continuous | – | °C |
| Explanatory | Wind speed (*Ws*) | Continuous | – | $m \cdot s^{-1}$ |
| Explanatory | Wind support (*Wp*) | Continuous | – | $m \cdot s^{-1}$ |
| Explanatory | Crosswind (*Wc*) | Continuous | – | $m \cdot s^{-1}$ |
| Explanatory | Thermal uplift | Continuous | – | $m \cdot s^{-1}$ |
| Explanatory | Age | Categorical | 2 (adult, 1to3-yo*) | – |
| Explanatory | Sex | Categorical | 2 (female, male) | – |
| Random | Bird ID | Factor | 35 | – |
| Random | Year | Factor | 4 | – |

* 1to3-yo = birds from 1 to 3 years old.

**Table S19** – Selected (ΔAIC < 7) generalized linear mixed models of environmental variables affecting the route choice in migrating White storks. First 10 models out of 39 are shown.

| Model | Variables | *w* | AICc | ΔAICc |
|---|---|---|---|---|
| 1 | Route choice × Wind support, Wind speed | 0.16 | 264.56 | 0.00 |
| 2 | Route choice × Wind support, Wind speed, Route choice × Age | 0.10 | 265.54 | 0.98 |

| | | | | |
|---|---|---|---|---|
| 3 | Route choice × Wind support, Wind speed, Route choice × Sex | 0.08 | 266.05 | 1.49 |
| 4 | Route choice × Wind support, Wind speed, Age | 0.06 | 266.45 | 1.90 |
| 5 | Route choice × Wind support, Route choice × Wind speed | 0.06 | 266.58 | 2.02 |
| 6 | Route choice × Wind support, Wind speed, Thermal uplift | 0.06 | 266.73 | 2.18 |
| 7 | Route choice × Wind support, Wind speed, Temperature | 0.04 | 267.15 | 2.60 |
| 8 | Route choice × Wind support, Route choice × Wind speed, Age | 0.04 | 267.58 | 3.02 |
| 9 | Route choice × Wind support, Wind speed, Sex | 0.03 | 267.76 | 3.20 |
| 10 | Route choice × Wind support, Route choice × Wind speed, Temperature | 0.02 | 268.46 | 3.90 |
| ... | ... | ... | ... | ... |

AIC: Akaike's information criterion, ΔAIC: difference in AIC between a given model and the best model, *w*: Akaike's weights.

**Table S20** – Model-averaged (subset models ΔAICc < 7) coefficients (*β*) with 95% confidence intervals (LCI, UCI) of generalized linear mixed models of environmental variables affecting the route choice in migrating White storks, ranked by their predictive importance (Σ*w*). *N* indicates the number of models containing the given variable.

| **Variable** | ***β*** | **LCI** | **UCI** | ***p*** | **Σ*w*** | ***N*** |
|---|---|---|---|---|---|---|
| (Intercept) | 9.04 | 8.35 | 9.74 | < 0.001 | – | – |
| Route choice | 1.01 | 0.26 | 1.77 | **0.008** | 1 | 39 |
| Wind support | –0.33 | –0.76 | 0.11 | 0.14 | 1 | 39 |
| Wind speed | 0.85 | 0.47 | 1.23 | **< 0.001** | 1 | 39 |
| Route choice × Wind support | –0.67 | –1.15 | –0.19 | **0.006** | 0.86 | 26 |
| Age | 0.27 | –0.71 | 1.25 | 0.59 | 0.30 | 12 |
| Route choice × Wind speed | –0.41 | –1.02 | 0.19 | 0.18 | 0.22 | 12 |
| Sex | 0.52 | –0.34 | 1.38 | 0.24 | 0.20 | 10 |
| Thermal uplift | 0.25 | –0.85 | 0.19 | 0.11 | 0.18 | 10 |
| Temperature | 0.18 | –0.11 | 0.48 | 0.21 | 0.14 | 8 |
| Route choice × Sex | –1.00 | –1.96 | –0.05 | 0.04 | 0.12 | 5 |
| Route choice × Age | 1.24 | –1.12 | 3.59 | 0.30 | 0.12 | 3 |
| Crosswind | 0.10 | –0.19 | 0.40 | 0.50 | 0.06 | 6 |
| Route choice × Thermal uplift | –0.33 | –0.85 | 0.19 | 0.21 | 0.02 | 1 |
| Route choice × Temperature | 0.31 | –0.15 | 0.76 | 0.19 | 0.02 | 1 |
| Route choice × Crosswind | 0.22 | –0.30 | 0.75 | 0.40 | 0.01 | 1 |

**Table S21** – Best model: Airspeed ~ Route_choice × Wind_support + Wind_speed + (1 | bird_id) + (1 | year) and information about the properties of its factors.

| **Variable** | ***β*** | **SE** | **t-value** | ***p*** |
|---|---|---|---|---|
| (Intercept) | 9.26 | 0.28 | 32.81 | < 0.001 |

| | | | | |
|---|---|---|---|---|
| Route choice | 1.04 | 0.28 | 3.69 | **< 0.001** |
| Wind support | –0.22 | 0.15 | –1.40 | 0.17 |
| Wind speed | 0.78 | 0.14 | 5.67 | **< 0.001** |
| Route choice × Wind support | –0.62 | 0.21 | –3.04 | **0.003** |

Random effects variance: Bird ID < 0.001, Year = 0.148

### Air speed – Fall – LMM

Full model formula: Air_speed ~ Route_choice × (Temperature + Wind_support + Crosswind + Wind_speed + Thermal_uplift + Ordinal_date + Sex + Age) + (1|Bird_ID) + (1|Year)

Number of obs. = 113

**Table S22** – Variables selected for the full model after checking for autocorrelation and collinearity of the most biologically meaningful variables.

| Model structure | Variable name | Variable type | N. levels (range or values) | Unit system |
|---|---|---|---|---|
| Dependent | Air speed | Continuous | _ | $m \cdot s^{-1}$ |
| Explanatory | Route choice | Binomial | 2 (0, 1) | – |
| Explanatory | Temperature | Continuous | – | °C |
| Explanatory | Wind support (*Wp*) | Continuous | – | $m \cdot s^{-1}$ |
| Explanatory | Crosswind (*Wc*) | Continuous | – | $m \cdot s^{-1}$ |
| Explanatory | Wind speed (*Ws*) | Continuous | – | $m \cdot s^{-1}$ |
| Explanatory | Thermal uplift | Continuous | – | $m \cdot s^{-1}$ |
| Explanatory | Ordinal date | Continuous | – | – |
| Explanatory | Age | Categorical | 2 (adult, 1to3-yo*) | – |
| Explanatory | Sex | Categorical | 2 (female, male) | – |
| Random | Bird ID | Factor | 67 | – |
| Random | Year | Factor | 5 | – |

* 1to3-yo = birds from 1 to 3 years old.

**Table S23** – Selected (ΔAIC < 7) generalized linear mixed models of environmental variables affecting the route choice in migrating White storks. First 10 models out of 40 are shown.

| Model | Variables | *w* | AICc | ΔAICc |
|---|---|---|---|---|
| 1 | Route choice, Wind support, Crosswind, Wind speed, Thermal uplift, Age | 0.15 | 306.90 | 0.00 |
| 2 | Route choice × Wind speed, Wind support, Crosswind, Thermal uplift, Age | 0.09 | 308.07 | 1.17 |
| 3 | Route choice, Wind support, Wind speed, Thermal uplift, Age | 0.08 | 308.17 | 1.26 |
| 4 | Route choice, Wind support, Crosswind, Wind speed, Thermal uplift | 0.08 | 308.29 | 1.39 |

| | | | | |
|---|---|---|---|---|
| 5 | Route choice, Wind support, Crosswind, Wind speed, Age | 0.07 | 308.43 | 1.53 |
| 6 | Route choice, Wind support, Crosswind, Wind speed | 0.07 | 308.61 | 1.70 |
| 7 | Route choice × Wind support, Wind speed, Thermal uplift, Age | 0.05 | 308.99 | 2.08 |
| 8 | Route choice × Age, Wind support, Wind speed, Thermal uplift | 0.03 | 309.89 | 2.99 |
| 9 | Route choice × Thermal uplift, Wind support, Wind speed, Thermal uplift | 0.03 | 310.13 | 3.22 |
| 10 | Route choice × Age, Wind support, Crosswind, Wind speed | 0.03 | 310.42 | 3.52 |
| ... | ... | ... | ... | ... |

AIC: Akaike's information criterion, ΔAIC: difference in AIC between a given model and the best model, *w*: Akaike's weights.

**Table S24** – Model-averaged (subset models ΔAICc < 7) coefficients ($\beta$) with 95% confidence intervals (LCI, UCI) of generalized linear mixed models of environmental variables affecting the route choice in migrating White storks, ranked by their predictive importance (Σ*w*). *N* indicates the number of models containing the given variable.

| **Variable** | $\beta$ | **LCI** | **UCI** | ***p*** | **Σ*w*** | ***N*** |
|---|---|---|---|---|---|---|
| (Intercept) | 8.83 | 8.48 | 9.17 | < 0.001 | – | – |
| Route choice | 0.75 | 0.31 | 1.18 | **< 0.001** | 1 | 40 |
| Wind support | –0.83 | –1.09 | –0.57 | **< 0.001** | 1 | 40 |
| Wind speed | 0.64 | 0.40 | 0.88 | **< 0.001** | 1 | 40 |
| Thermal uplift | 0.35 | 0.09 | 0.61 | **0.007** | 0.72 | 24 |
| Age | –0.43 | –0.81 | –0.05 | **0.027** | 0.63 | 17 |
| Crosswind | 0.55 | –0.55 | –0.08 | **< 0.001** | 0.61 | 25 |
| Route choice × Wind speed | –0.25 | –0.73 | 0.22 | 0.29 | 0.13 | 5 |
| Route choice × Wind support | –0.24 | –0.64 | 0.14 | 0.22 | 0.11 | 6 |
| Route choice × Age | –0.31 | –1.06 | 0.44 | 0.41 | 0.06 | 2 |
| Sex | –0.13 | –0.53 | 0.26 | 0.51 | 0.06 | 6 |
| Route choice × Thermal uplift | –0.23 | –0.58 | 0.13 | 0.21 | 0.06 | 3 |
| Route choice × Crosswind | 0.13 | –0.35 | 0.61 | 0.60 | 0.04 | 3 |
| Ordinal date | 0.11 | –0.09 | 0.31 | 0.30 | 0.04 | 6 |
| Temperature | –0.04 | –0.27 | 0.19 | 0.74 | 0.03 | 4 |
| Route choice × Sex | 0.58 | –0.10 | 1.26 | 0.088 | 0.10 | 2 |

**Table S25** – Best model: Air_speed ~ Route_choice + Wind_support + Wind_speed + Thermal_uplift + Age + Crosswind + (1 | bird_id) + (1 | year) and information about the properties of its factors.

| **Variable** | $\beta$ | **SE** | **t-value** | ***p*** |
|---|---|---|---|---|
| (Intercept) | 8.84 | 0.15 | 58.23 | < 0.001 |

| | | | | |
|---|---|---|---|---|
| Route choice | 0.81 | 0.17 | 4.05 | < **0.001** |
| Wind support | –0.85 | 0.11 | –7.91 | < **0.001** |
| Thermal uplift | 0.28 | 0.11 | 2.66 | **0.009** |
| Crosswind | –0.25 | 0.09 | –2.62 | **0.01** |
| Wind speed | 0.61 | 0.11 | 5.72 | < **0.001** |
| Age | –0.40 | 0.18 | –2.24 | **0.027** |

Random effects variance: Bird ID = 0.056, Year = 0.021

**Table S26** – Summary table of best models

| **Outcome** | **Season** | **Variable** | ***β*** | **SE** | ***t*** | ***p*** |
|---|---|---|---|---|---|---|
| Ground speed | Spring | (Intercept) | 9.19 | 0.28 | 32.42 | < 0.001 |
| | | Route choice | 0.54 | 0.25 | 2.13 | **0.035** |
| | | Wind support | 0.98 | 0.18 | 5.41 | < **0.001** |
| | | Route choice × Wind support | –0.78 | 0.24 | –3.18 | **0.002** |
| | Autumn | (Intercept) | 9.96 | 0.15 | 66.47 | < 0.001 |
| | | Route choice | 0.74 | 0.20 | 3.62 | < **0.001** |
| | | Wind speed | 0.36 | 0.09 | 3.73 | < **0.001** |
| | | Thermal uplift | 0.37 | 0.10 | 3.61 | < **0.001** |
| | | Age | –0.49 | 0.19 | –2.60 | **0.011** |
| Air speed | Spring | (Intercept) | 9.26 | 0.28 | 32.81 | < 0.001 |
| | | Route choice | 1.04 | 0.28 | 3.69 | < **0.001** |
| | | Wind support | –0.22 | 0.15 | –1.40 | 0.17 |
| | | Wind speed | 0.78 | 0.14 | 5.67 | < **0.001** |
| | | Route choice × Wind support | –0.62 | 0.21 | –3.04 | **0.003** |
| | Autumn | (Intercept) | 8.84 | 0.15 | 58.23 | < 0.001 |
| | | Route choice | 0.81 | 0.17 | 4.05 | < **0.001** |
| | | Wind support | –0.85 | 0.11 | –7.91 | < **0.001** |
| | | Thermal uplift | 0.28 | 0.11 | 2.66 | **0.009** |
| | | Crosswind | –0.25 | 0.09 | –2.62 | **0.01** |
| | | Wind speed | 0.61 | 0.11 | 5.72 | < **0.001** |
| | | Age | –0.40 | 0.18 | –2.24 | **0.027** |

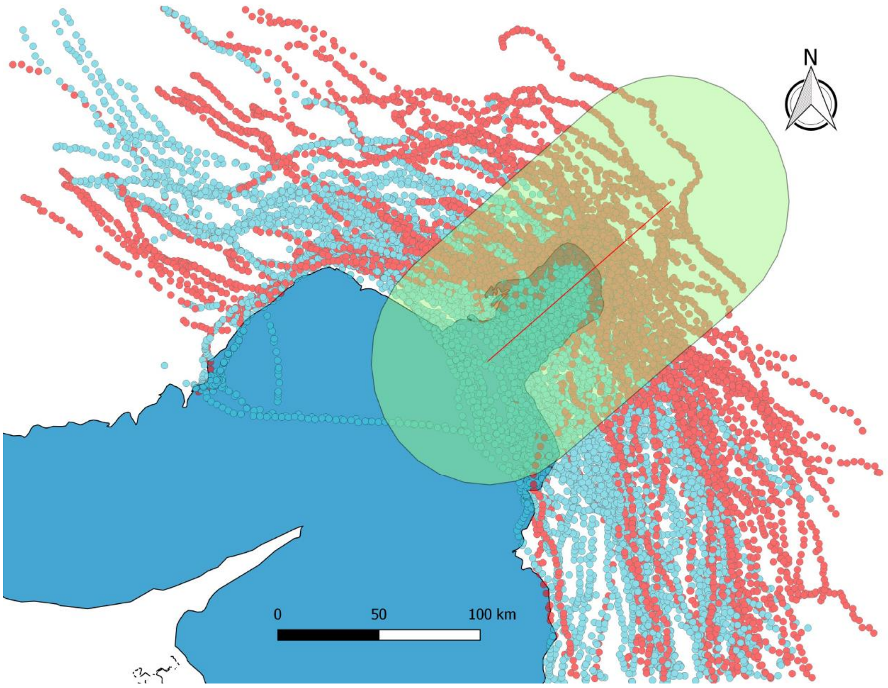


**Figure S18** – Illustration of the Iskenderun bay area and GPS locations of the white storks. The selected points included around the bay by the buffer of 60 km around the line dividing the bay in half are used to illustrate the differences in flight time and distance covered in a defined space around the bay including before and after the sea crossing or land detour (see Figure S19).

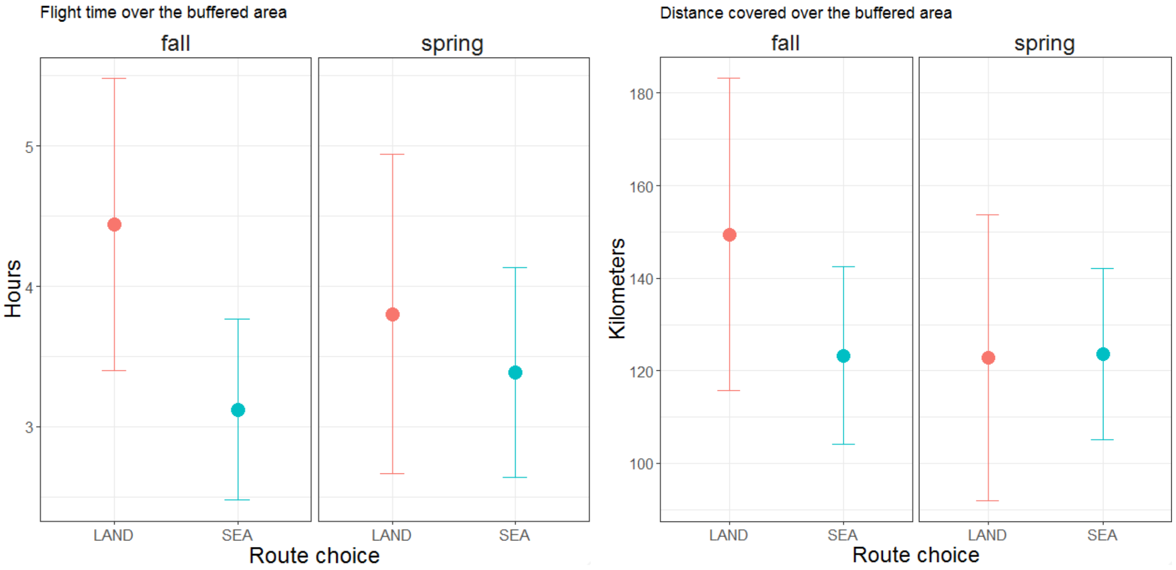


**Figure S19** – Flight time (left) and distance covered (right) by route choice (LAND vs. SEA) of white storks in the different seasons relative to the buffer area in Figure S18.

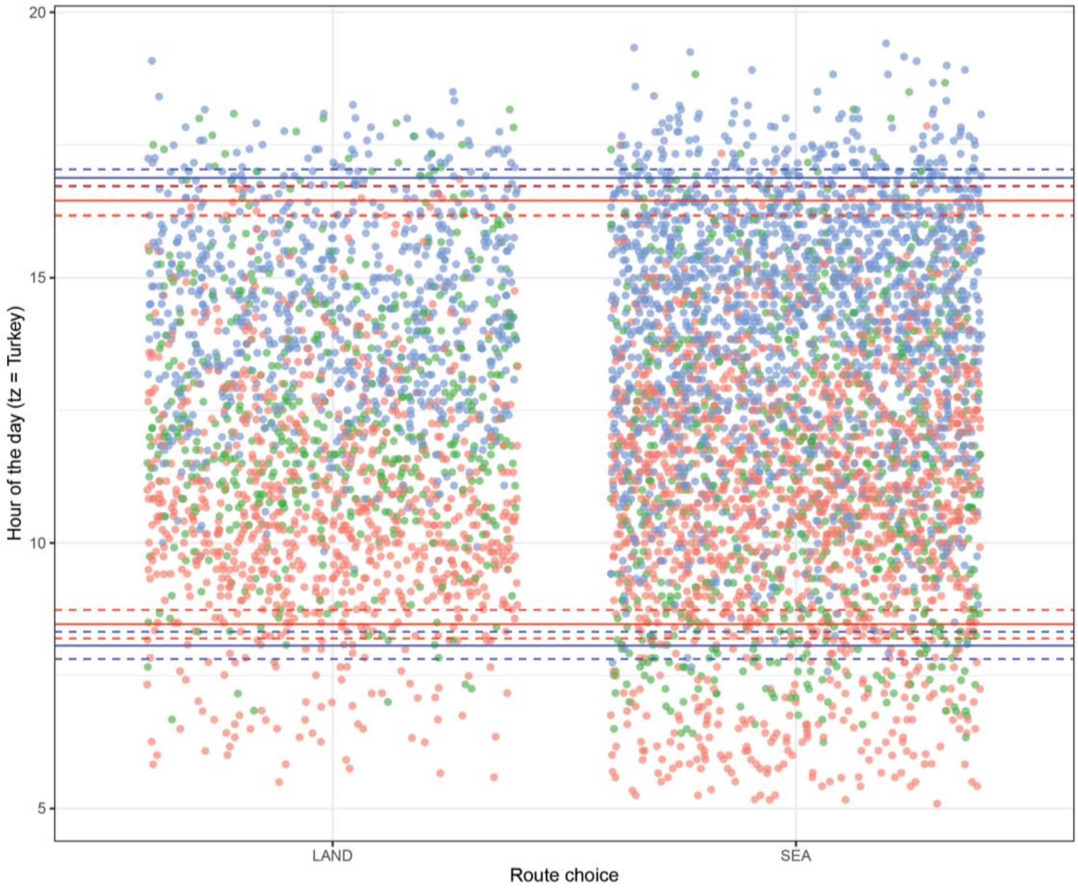


**Figure S20** – White storks locations in spring in relation to the local time grouped by route choice. Colours show the section in relation to the Iskenderun Bay where the locations were recorded: BEFORE (red), ACROSS (green) and AFTER the bay (blue). Continuous lines are mean ± S.E. (dashed lines) of starting and ending time of the daily migration trip over the area, in blue for the group that crossed the bay and in red for the detour.

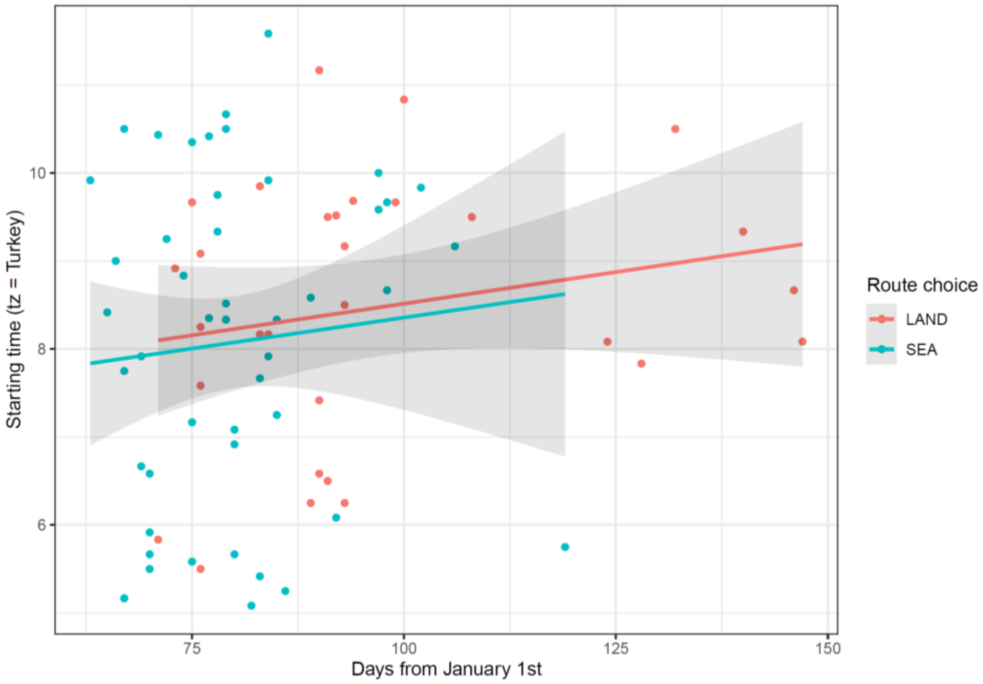

**Figure S21** – Relationship between the time of daily migration onset and ordinal date of white storks during spring migration (starting time is relative to the migration day over or around the Iskenderun Bay).

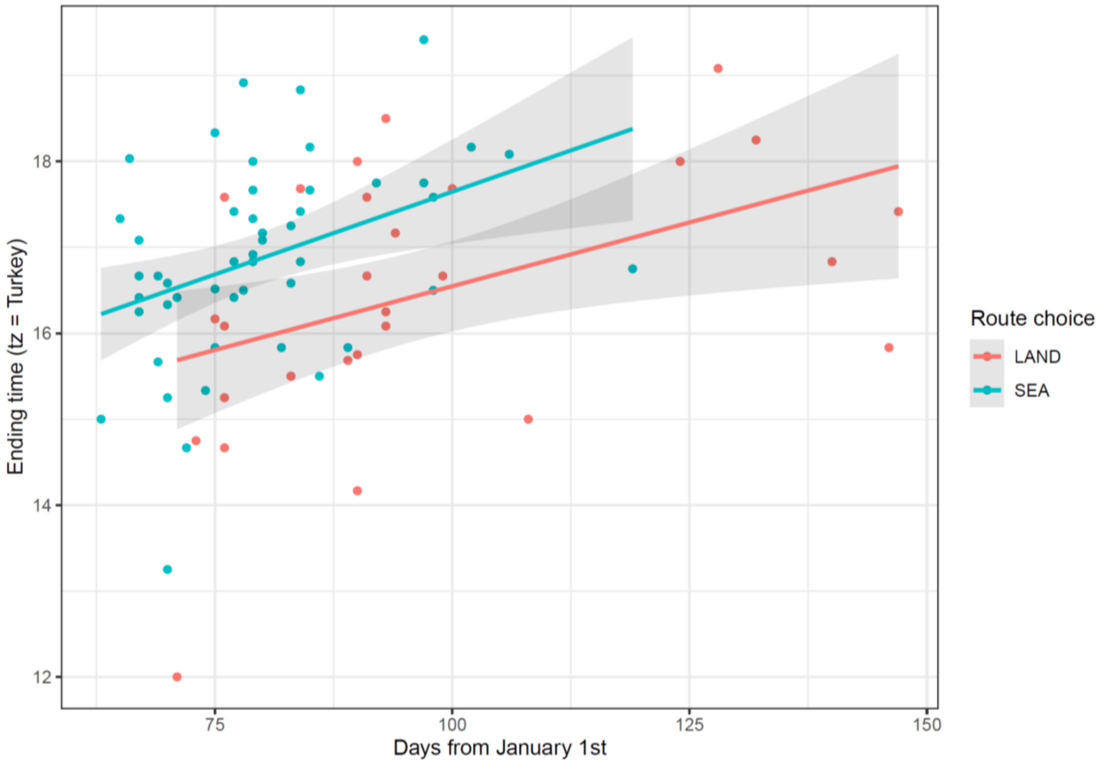


**Figure S22** – Relationship between the time of daily migration end and ordinal date of white storks during spring migration (ending time is relative to the migration day over or around the Iskenderun Bay).

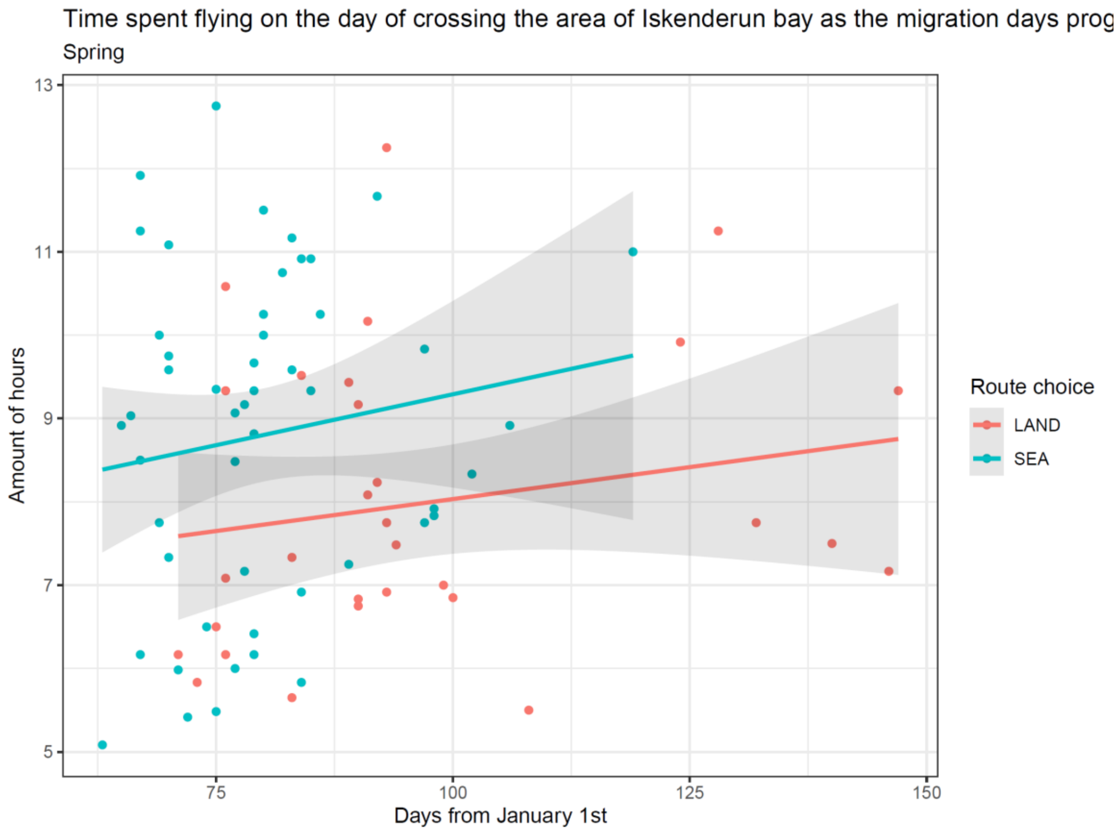

**Figure S23** – Relationship between the daily migrating duration and ordinal date of white storks during spring migration (amount of time is relative to the migration day over or around the Iskenderun Bay).
